# Component specific responses of the microbiomes to common chemical stressors in the human food chain

**DOI:** 10.1101/2024.04.20.590402

**Authors:** Wasimuddin, Aurea Chiaia-Hernandez, Céline Terrettaz, Lisa Thoenen, Veronica Caggìa, Pierre Matteo, Miquel Coll-Crespi, Matheus Notter, Mohana Mukherjee, Teresa Chavez-Capilla, Francesca Ronchi, Stephanie C. Ganal-Vonarburg, Martin Grosjean, Moritz Bigalke, Sandra Spielvogel, Andrew Macpherson, Adrien Mestrot, Siegfried Hapfelmeier, Matthias Erb, Klaus Schlaeppi, Alban Ramette

## Abstract

Along a food chain, microbiomes occur in each component and often contribute to the functioning or the health of their host or environment. ‘One Health’ emphasizes the connectivity of each component’s health. Chemical stress typically causes dysbiotic microbiomes, but it remains unclear whether chemical stressors consistently affect the microbiomes along food chain components. Here, we systematically challenged a model food chain, including water, sediments, soil, plants, and animals, with three chemical stresses consisting of arsenic (a toxic trace element), benzoxazinoids (an abundant bioactive plant metabolites), and terbuthylazine (an herbicide typically found along a human food chain). The analysis of 1,064 microbiome profiles for commonalities and differences in their stress responses indicated that chemical stressors decreased microbiome diversity in soil and animal, but not in the other microbiomes. In response to stress, all food chain communities strongly shifted in their composition, generally becoming compositionally more similar to each other. In addition, we observed stochastic effects in host-associated communities (plant, animal). Dysbiotic microbiomes were characterized by different sets of bacteria, which responded specifically to the three chemical stressors. Microbial co-occurrence patterns significantly shifted with either decreased (water, sediment, plant, animal) or increased (soil) network sparsity and numbers of keystone taxa following stress treatments. This suggested major re-distribution of the roles that specific taxa may have, with the community stability of plant and animal microbiomes being the most affected by chemical stresses. Overall, we observed stress- and component-specific responses to chemical stressors in microbiomes along the model food chain, which could have implications on food chain health.

## Introduction

The ‘One Health’ concept emphasizes the ecological relationships and interdependencies of all components of a system to collectively determine the global health of that system^1^. Hence, the health of our planet results from a connected health of us, plants, animals and the environment. All system components host microbial communities or are colonized by microbiomes that have important roles in the health of each system component. The One Health concept was extended to include the full breadth of microbiomes^2–4^. It is thought that a microbiome perspective strengthens the One Health concept due to i) the contribution of microbiomes to the health of individual system components, ii) the importance of microbiome processes for the transfer of energy, matter and chemicals between system components, and iii) the vital services provided by microbiomes to overall system’s health. Furthermore, dysbiotic microbiomes of humans^5^, plants^6^, animals^7^ or the environment^8^ are often associated with diseases or impaired ecosystem performance. A dysbiotic state can originate from stressors that either induce deterministic, stochastic, or a mix of these two, effects on the microbiome and thereby reduce the ability of the host or its microbiome to regulate community composition^9^. What is not well studied in One Health context is whether stressors of a whole system influence the microbiomes of the diverse components with commonalities and/or disparities in their responses. Such information is crucial for estimating individual component microbiomes’ resilience against common disturbances in a One Health framework.

A wide range of common environmental and anthropogenic stressors including chemicals like toxic trace elements^10^, bioactive plant metabolites^11^, and pesticides^12^ are known to negatively affect the health of different system components. Such stresses can directly impact the health of the exposed environment or organism, for instance by changing metabolic rates, inhibiting enzymatic functions or indirectly via perturbating or throwing off balance the microbiome’s composition. Research has traditionally focused on understanding the direct stress effects on host or environmental physiology (toxic trace elements^13^, plant metabolites^14^, and pesticides^15,16^) as well as direct stress effects on microbiomes of individual system components. For instance, water, soil, plant or animal associated microbiomes are perturbed by stresses like toxic trace elements, plant metabolites, and pesticides^17–19^. However, the indirect and microbiome-mediated contributions to connected system components, i.e. taking the One Health perspective, have received much less attention. A major gap towards such One Health understanding, is the lack of systematic studies where microbiomes of different system components are challenged with the same stressors, at the same doses and with the same exposure protocol. Such systematic work will allow to specifically answer fundamental questions of a One Health framework, such as (i) whether microbiomes of diverse system components can be perturbed with the same stress exposure protocol, (ii) if yes, how does their stress sensitivity compares in direction or magnitude within and across components, and (iii) whether there are commonalities or differences in the microbiomes’ stress responses from different system components?

To close this gap, we set up an idealized food chain system represented by water and sediment, soil, plants and animals (**Figure 1A**). Our experimental food chain consisted of environmental components (water, sediment and soil) with free-living microbial communities of high diversity, of primary producers (plants, i.e. corn root microbiomes), and end-consumers (animals, i.e. mouse gut microbiomes) with low-diversity host-associated microbiomes^20^. For systematic challenging of these different system components, we selected three chemical stressors found along a food chain that are known to impact human and/or environmental health: Arsenic (As) is a toxic trace element, Benzoxazinoids (Bx) are bioactive plant metabolites, and terbuthylazine (Tb) is a potent herbicide.

**Figure 1.**
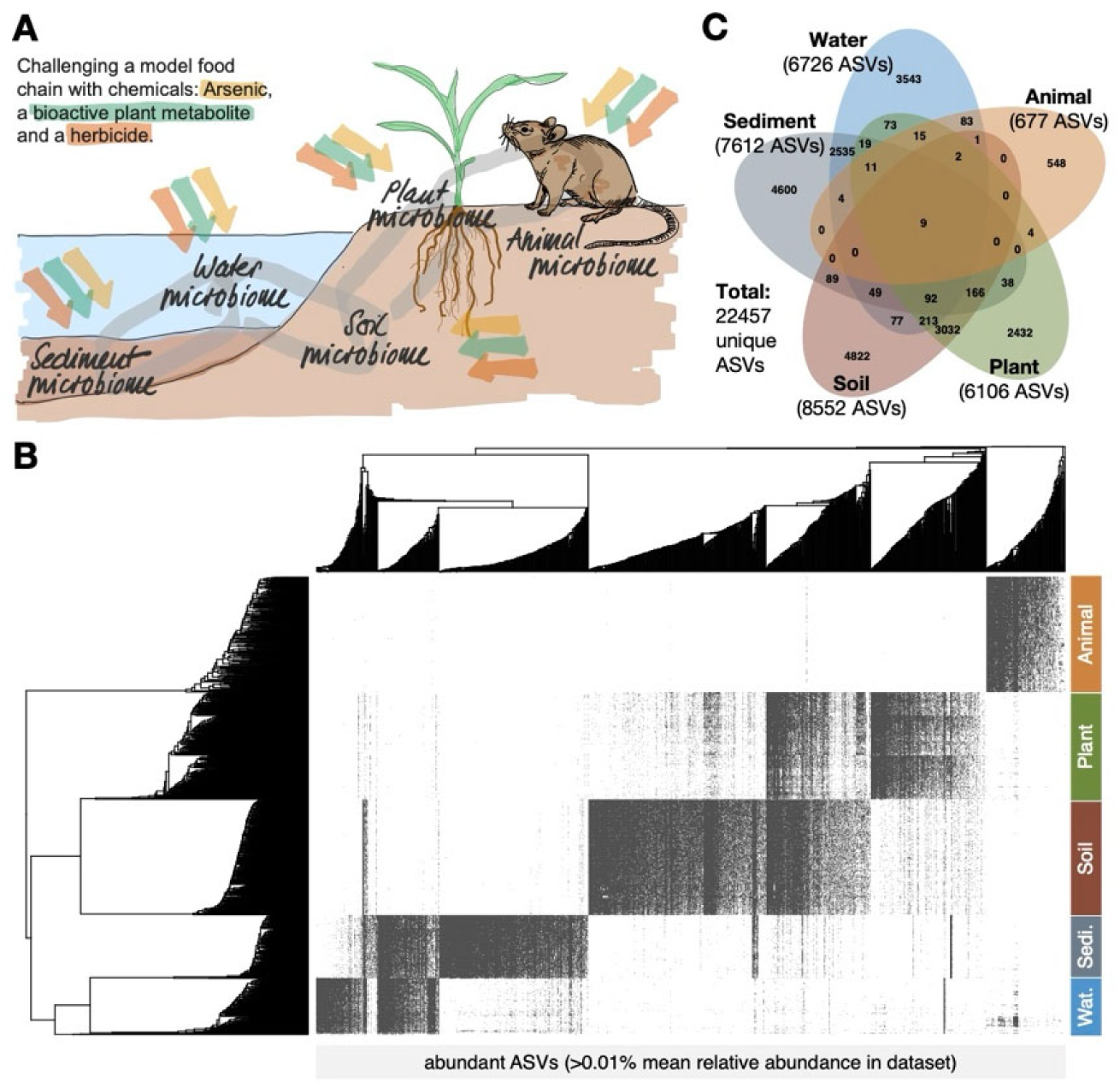
**A**) Depiction of representative components of a model food chain used in this study. **B**) Heatmap of abundant ASVs (>0.01% mean relative abundance in the dataset) across components (right) showing little overlap among components except between related components (water and sediment, soil and plant), arranged according to hierarchical cluster tree. **C**) Venn diagram showing unique and shared ASVs among components.

The element As is ubiquitously found in many environments and functions as a carcinogen for humans^21^. Contamination by As presents a global catastrophe with around 1.5% of the world’s population suffering from As exposure through drinking water^22,23^, rice^24^ or corn consumption^25^. Although high levels of naturally occurring As can be found in groundwater and soils around the globe, it is most problematic in countries with dense population and lack of infrastructure to detect and manage As contamination. As is known to impact microbiomes in some of the tested system components^26,27^, but its influence on diverse microbiomes of a food chain is not known. We utilized inorganic As^V^ for this study as this is the most abundant form of As found in the environment^28^.

Bx are probably the most relevant plant secondary metabolites in food chains. This is because they are highly abundant in agroecosystems as they are secreted in large quantities to soil by *Poaceae* plants (sweet grasses) that include the widely grown crops like corn, wheat and rye^29^. Secondary metabolites generally play vital roles in plant adaptation to the environment^30^. Bx are a group of highly bioactive multifunctional compounds that act as feeding toxins against herbivores^31^, have antimicrobial activities against microbes^32^ and improve plant nutrient acquisition^33^. Bx could have direct influence on health of diverse food chain components including humans^34^. Bx were found to affect microbiomes as Bx-conditioned soils mediated growth and defence effects on the following plant generation^35^, forwarding Bx as a chemical for One Health research. Whether Bx influence other microbiomes than those of plants and soil is currently unknown. We choose the Bx 6-methoxy-benzoxazolin-2-one (MBOA) for this study, because this compound is stable and abundantly accumulates in soils^36^.

Tb is a broad-spectrum herbicide from the chloro-*s*-triazine group, which is commonly used for chemical weed control around the globe^37^ and has been detected in different environments^38,39^. More generally, pesticides comprising herbicides, insecticides, and fungicides are not only used in agroecosystems, but also in other areas to protect humans from various pests and diseases. Besides their specific toxicity against weeds, pest insects and pathogenic fungi, they also cause many negative health and environmental side-effects on off-target organisms^40^. Pesticides are associated with significant morbidities and mortalities each year^41^. They are also known to influence microbiomes but their relative impact on diverse food chain microbiomes is not known^42^. We included Tb in this study, because it is broadly used, and because the compound and its degradation products have been found in surface and groundwater^38^ and accumulate in soils^43^ and sediments^39^, displaying long-term stability in the environment^43^.

Finally, in large-scale experiments, we systematically challenged the microbiomes of the different food chain components with As, Bx and Tb. For this, we developed specific setups and application procedures for continuous exposure of the microbiome to the chemicals (**Figure S1**). To assess their sensitivity to chemical perturbation, we exposed the microbiomes to the same concentrations and sampled them following the same timeframe. The overall aim of the experiments was to identify the microbiomes that are the most resistant, and conversely those that are the most sensitive ones to the chemical stresses, i.e. to find the Achilles’ heel of our experimental food chain.

## Materials and Methods

### Experimental overview

We studied the components of our idealized food chain consisting of water, sediment, soil, plant and animal in three large, parallelly executed experiments (**Figure S1**; termed the ‘water/sediment’, the ‘soil/plant’ and the ‘animal’ experiments). We developed specific application procedures to ensure that the microbiomes in the different components are continuously exposed to the same chemical concentrations. In the water/sediment and soil/plant experiments, we applied the chemicals in daily intervals to approximate continuous exposure, whereas this was achieved through drinking water in the animal experiment. Besides continuous exposure, the application procedures were conceived that the chemicals reached 1x final concentrations of 10, 100 and 1,000 µg/L in each system component. In the **Supplementary Methods** we provide the experimental details related to sources, setup, chemical application procedures and sampling of the three experiments. After application of the chemicals, we collected samples from each component for microbiome analysis at two time points, 1 (d1) and 7 (d7) days post application.

### Treatment solutions of the chemicals

We prepared the “treatment solutions” of the chemicals As (Sodium arsenate dibasic heptahydrate, ≥98% purity; Sigma-Aldrich, Germany), Bx (6-methoxy-benzoxazolin-2-one (MBOA), >98% purity; Sigma-Aldrich) and Tb (grade analytical grade; Pestanal, Germany) with specific concentrations for the different experiments. The water/sediment experiments needed highly concentrated treatment solutions (300x, to minimize dilution by the lake water and its microbiome), while we prepared 3x and 1x treatment solutions for the soil/plant and the animal experiments, respectively (**Figure S1**). The treatment solutions of As and Bx were prepared in water, which was sufficient as a buffer as the added compounds did not change the pH of the solutions (data not shown). Because the animal experiment required ultrapure surgical irrigation water (ERKF7114; Baxter, USA), we used this water as common source to prepare all treatment solutions. Tb, however, was dissolved in pure ethanol (>98% purity, Sigma-Aldrich, Germany) due to its insolubility in water. Ethanol was also added to control treatment solution for Tb at final amounts of 0.3%.

### DNA Extraction, 16S rRNA amplicon library preparation and sequencing

DNA was extracted using the DNeasy PowerSoil HTP 96 Kit from Qiagen (Hilden, Germany), as recommended by the Earth Microbiome Project EMP^20^ following the manufacturer’s protocol. Loading of the sample material to the DNA extraction plates was done as follows: Water samples were pipetted directly on to plates (250 µL, well-homogenized by vortexing). Sediment samples were defrosted, briefly vortexed, centrifuged, and pipetted up and down for homogenization of water and solid particles, then 250 µL was transferred into the extraction plates. A sterile spatula was used to retrieve 230-250 mg of defrosted soil samples. Corn roots were lyophilized for 48 h, and after grinding, 15 mg of fine ground powder was then used to load the plate. The loading unit for mouse samples corresponded to one to two faecal pellets. For loading the extraction plates, we processed the samples in batches by sample type to avoid cross-contamination and because of the different handling units (weight, unit or volume). Within batches, the sample groups (time point and treatment) were randomly positioned on the plate. With this randomization scheme, we tackled the practical challenges (diverse input materials, avoiding sample mix-up) without compromising scientifically rigorous treatment comparisons (randomization of treatment groups). DNA was eluted in 50 µL (water, sediment), 75 µL (plant) or 100 µL (soil, animal) of C6 buffer of the kit (10 mM Tris-Cl, no EDTA) and stored at -20°C until further use.

We then performed a bulk adjustment of DNA concentrations for the water, sediment, soil, plant and animal samples as follows: From each sample type, we measured the DNA concentrations of 20 random samples from different DNA extraction plates using Nanodrop (Thermo Fischer, Waltham, USA). The average DNA concentration of a sample type was taken to bulk-adjust all samples of the same sample type. PCR-ready concentrations were set at 10 ng/µL (sediment, soil and plant samples) and 1 ng/µL (animal samples). DNA concentrations of water extracts were <1 ng/µL and were used without further dilution. At this step, samples were re-organized for amplification and assigned to 5 different sequencing libraries (L1 to L5). Each library consisted of ∼240 samples, with replicates of a treatment group being present in at least two different sequencing libraries.

Bacterial 16S rRNA gene amplicon libraries were prepared using PCR primers, reagents and cycling conditions as recommended by the EMP^20^, according to our previous study^44^: We barcoded the amplicons with the Access Array barcode system from Fluidigm in a two-step approach adapted from Illumina’s standard 16S profiling protocol. In the first step, we performed target gene (16S rRNA, region V4) amplification using the PCR primers 515F and 806R^45,46^ coupled to CS1 and CS2 linker sequences (CS1-515F: 5’- ACACTGACGACATGGTTCTACA-GTGYCAGCMGCCGCGGTAA-3’ and CS2-806R: 5’- TACGGTAGCAGAGACTTGGTCT-GGACTACNVGGGTWTCTAAT-3’) of the Access Array barcode system, respectively. PCR reactions (20 µL total volume) were prepared in a UV-irradiated PCR hood, and contained 0.8x Platinum Hot Start PCR Master Mix (Thermo Fisher), 0.2 µM of each primer, PCR-grade water and 3 µL of DNA template. After 3 min initial denaturation at 94°C, we ran 25 PCR cycles (25 of the 35 cycles as suggested by EMP^20^; 45 s at 94°C, 60 s at 50°C and 90 s at 72°C) followed by 10 min final elongation at 72°C. We performed gel electrophoresis with few samples, and the positive and negative controls for each PCR plate to confirm that the PCR has worked and was not contaminated. PCR products were purified with self-made Solid Phase Reversible Immobilisation (SPRI) magnetic beads (https://openwetware.org/wiki/SPRI_bead_mix).

In the second PCR step, we barcoded the individual samples with the Access Array system consisting of 384 barcodes (BC). The PCR primers PE1-CS1-F and PE2-[BC]-CS2-R contain the paired-end (PE) adapters required for Illumina sequencing and bind via the linker sequences CS1 and CS2 to the PCR amplicons of the first step. Stocks (50 µL, 2 µM) of the 384 unique primer combinations were repeatedly utilized to prepare the 5 libraries L1 to L5. PCR reactions were prepared in volumes of 25 µL with 0.8x Platinum Hot Start PCR Master Mix (Thermo Fisher Scientific, Reinach, Switzerland), Access Array primers (0.4 µM), PCR-grade water and 5 µL of the purified PCR product as template. After 3 min of initial denaturation at 94°C, we ran 10 PCR cycles (25 cycles in step 1 + 10 cycles in step 2 correspond to the 35 cycles as suggested by EMP^20^; 45 s at 94°C, 60 s at 60°C and 90 s at 72°C), followed by 10 min of final elongation at 72°C. Again, gel electrophoresis was performed with few samples, and positive and negative controls for each PCR plate to confirm that the PCR has worked. No DNA contamination was observed in the negative controls after two rounds of PCR amplification. Amplicon DNA of the second PCR was purified with SPRI beads as described above and quantified with NanoDrop 8000 (Thermo Fisher Scientific).

For equimolar pooling of the barcoded amplicons into their assigned library (L1 to L5), we used a robotic liquid handling station (Brand, Wertheim, Germany). Pooled libraries were well mixed and a subset was purified using the SPRI beads as described above. DNA concentration and size of the purified library were then determined by Qubit 1.0 (Thermo Fischer) and TapeStation (Agilent, Santa Clara, CA, USA) analyses. The final pooled libraries were paired-end sequenced (2 × 300 cycles) in five runs on Illumina MiSeq at the NGS platform of University of Bern (www.ngs.unibe.ch). The sequencing data is available from the European Nucleotide Archive (http://www.ebi.ac.uk/ena) under the study accession PRJEB72104.

### Bioinformatic and statistical analyses

All code and metadata (experimental design, sample-to-barcode assignments) are available on GitHub (https://github.com/wasimbt/Component-specific-responses-of-the-microbiome). Demultiplexed reads without barcodes and adapters were received as output from the sequencing centre. Primer sequences were removed by using *Cutadapt*^47^ (version 2.5). All subsequent analyses were performed within the R environment^48^ (version 3.5.1). For data pre-processing, we followed the *DADA2*^49^ pipeline (version 1.10.1) by keeping the same parameters for the five libraries, except for error rate estimation that was allowed to be library-specific. Reads were trimmed from both ends based on quality profile, error rates were learned from the data using the parametric error model as implemented in *DADA2*. After denoising and merging of forward and reverse reads, all five libraries were regrouped. Chimeric sequences were removed from the dataset by following the ‘consensus’ method implemented in *DADA2.* The final table thus consisted of number of occurrences of amplicon sequence variants (ASVs; i.e. sequence groups differing by as little as one nucleotide) in each sample. Taxonomy assignments of the ASVs were performed using the naïve Bayesian classifier^50^ with the SILVA database^51^ (version v132, non-redundant). Species level assignment was done by exact matching (100% identity) of ASVs with database sequences, as previously recommended. *Phyloseq*^52^ (version 1.24.2) was used for further data processing. We removed ASVs with less than 10 read counts from overall dataset. Furthermore, ASVs belonging to chloroplast, mitochondria, and unassigned ASVs at phylum level were removed from the dataset.

*Alpha and beta diversity analyses:* We investigated the effects of sample type (water, sediment, soil, plant, animal), treatments (Control (Ctr), As, Bx, Tb), time point (0, 1, 7 days), concentration (0, 10, 100, 1,000 µg/L), and interactions among these factors. First, we analysed bacterial diversity for each sample using two different alpha diversity indices (number of observed species and Shannon) after rarefying the data to 8,100 sequences per sample using *Phyloseq*^52^. To analyse the effects of these factors on alpha diversity, we performed General Linear Modelling (GLM) by using the *lme4*^53^ package (version 1.1.30). As the samples were pooled in 5 libraries and were sequenced in five sequencing runs, we also included “library identity” as explanatory factor in the model to account for potential technical confounding. We performed Tukey’s Honest Significant Difference test (HSD) to compare average effects between groups when overall multivariable model significance was observed. Second, beta diversity analyses were calculated based on a Bray-Curtis dissimilarity matrix after rarefying the data to 8,100 sequences per sample using *Phyloseq*^52^. The permutational multivariate analysis of variance (PERMANOVA) was employed as implemented in the *adonis* function of the *vegan*^54^ package (version 2.5-2) to test the significance of the differences in community composition with 999 permutations. For beta diversity metric, we similarly included sample type, treatments, time point, concentration and interactions among these factors in the model as explanatory variables. We also included “library identity” as potential confounding factor in the model. We performed *pairwise.adonis* to compare groupings, similar to Tukey’s HSD done on linear models. To visualize patterns of separation between different sample groups, non-metric multidimensional scaling (NMDS; *Phyloseq*) plots were prepared based on Bray-Curtis dissimilarity matrices. To assess the strength of treatment in each specific component of the food chain, we performed constrained ordination (distance-based redundancy analysis; dbRDA) by using the *capscale* function of the *vegan*^54^ package on Bray-Curtis dissimilarity matrices within each component. We employed ANOVA to assess the significance of each component model.

In order to understand whether treatments reflect true shift in microbial community composition or differential spread (dispersion) of data points from their group centroid, we assessed the multivariate homogeneity of group dispersions by performing PERMDISP test using the *betadisper* function of the *vegan*^54^ package on Bray-Curtis dissimilarity matrices. We employed permutation (999) test with *permutest* function as implemented in *vegan*^54^ to analyse significance of grouping (treatment) for each component of the food chain.

*Differential abundance analyses:* In order to identify ASVs specifically influenced by a given treatment, we employed a negative binomial model-based approach available in the *DESeq2*^55^ package (version 1.22.2), in which ASV relative abundances were compared for each treatment vs. control group (Ctr-As, Ctr-Bx, Ctr-Tb) for the respective component. Only ASVs remained significant (*P*≤0.05) after Benjamini–Hochberg correction of Wald test were considered as differently abundant ASVs. Here, we calculated an Influence Score (IS) for each comparison, which considers both the number of affected ASVs and their relative change in abundance, as consists in the cumulative log-fold changes for all ASVs significantly differing for a given comparison (e.g. Ctr-As; **Figure S6**).

*Network analyses:* To infer the relationships among ASVs, we prepared networks for each food chain component and their treatments by using Sparse Inverse Covariance Estimation for Ecological Association Inference SPIEC-EASI^56^. SPIEC-EASI is a statistical method for the inference of ecological networks that relies on algorithms for sparse neighbourhood and inverse covariance selection, and that applies data transformation and normalization, which can better deal with compositional data. To prepare the networks, only ASVs present in control groups were kept in treated groups for each component. Furthermore, ASVs containing fewer than 100 reads from overall component dataset, present in less than 15% of the control samples were removed prior to selecting control ASVs. Network inference used the Meinshausen-Buhlmann method for neighbourhood selection and the bounded StARS approach with nlambda of 50 and 99 pulsar permutations. Node attributes, such as degree distribution, betweenness centrality, transitivity, closeness centrality, were calculated using the *igraph*^57^ package (version 1.3.2) with 10,000 iterations. We then performed the Kolmogorov–Smirnov test to compare node attributes between control and treated groups. Kolmogorov–Smirnov test compares the overall shape of the cumulative distribution of two variables where the null hypothesis is that the variables derive from the same distribution. To characterize the underlying network degree distribution type, we evaluated four distributions namely, power-law, log normal, exponential and Poisson and tested goodness of fit of the distribution after 1,000 iterations, and we also compared the fitted distributions with each other’s using *t*-test to detect the best fitting distribution(s). Finally, to detect hub nodes which could represent keystone taxa, we calculated Kleinberg’s hub centrality scores using the *hub_score* function implemented in *igraph*^57^. Nodes having hub score values of more than 0.7 were assigned as hub nodes across sample types.

## Results

### Microbiomes are distinct across components of a food chain

To assess the sensitivity of different food chain components to chemical perturbation, we systematically exposed their microbiomes to As, Bx and Tb concentrations of 10, 100 and 1,000 µg/L and sampled them after same exposure times (**Figure S1**). Each food chain component was also treated with buffer and these control microbiomes served as baseline for healthy un-perturbed microbiomes. We first validated the quantities of As, Bx and Tb in the water or treatment solutions, which we used to challenge the different food chain microbiomes. The water of the water/sediment and animal experiments and the treatment solutions of the soil/plant experiment contained the standardized 10-fold increments of stress chemicals at the expected concentrations (**Figure S2**). Following the systematic exposures, we collected samples from each component at two time points, and thus characterized a total of 1,064 microbiomes originating from water (n= 133; Ctr-34, As-33, Bx-31, Tb-35), sediment (n= 144; Ctr-36, As-36, Bx-36, Tb-36), soil (n= 266; Ctr-36, As-78, Bx-77, Tb-75), plants (n= 255; Ctr-37, As-72, Bx-75, Tb-71) and animal (n= 266; Ctr-25, As-82, Bx-76, Tb-83). All samples were subjected to high-throughput sequencing of the V4 region of the bacterial 16S ribosomal RNA gene. We recovered on average 38’709 (range 5’437-116’475) high-quality, taxonomically assigned reads per sample. Microbiome diversity differed markedly between the different food chain components (**Figure S4**). Quantifying their effect size from PERMANOVA (based on interpreting the R^2^ of the model) revealed 71.8% based on the Bray-Curtis metric (**Table S2**). The ‘naturally close’ microbiomes of water and sediment, as well as of soil and plant each shared some abundant bacteria, whereas hardly any overlap existed between these microbiomes and the distinct animal microbiomes (**Figure 1B**). The same was true when inspecting all ASVs that were detected in this study (**Figure 1C**). Hence, it is unlikely that the stress treatments will affect the different bacteria of the food chain components in similar manner. Thus, we further analysed the stress-induced impact on the microbiomes of the different components separately, at the levels of diversity patterns, community composition and interaction between members of the community.

### Chemical stressors mainly affect diversity of soil and animal microbiomes

We first investigated to which degree the chemical stressors perturbed alpha diversity in food chain microbiomes. Sediment and soil harboured the richest (number of observed ASVs; **Figure 2A**) and most diverse (Shannon; **Figure S5A**) microbiomes followed by water and plant microbiomes, while animal microbiomes were lowest in both metrics. We used General Linear Modelling (GLM) to statistically assess and quantify the effects of the applied stressors (As, Bx, Tb) on bacterial richness when compared to un-perturbed conditions (**Table S3**). Richness was most strongly (interpreting the sums of squares of the model as effect size) differing between sample type, followed by time point, concentration and type of chemical treatment, while accounting for technical variation due to sequencing library preparation (all *P*<0.001). Albeit of lower effect size, many factor interactions including chemical treatment and concentrations were also significant. Significant differences on richness were observed between control and treatment groups for soil microbiomes, where chemical stressors reduced richness (**Figure 2B**). No chemical stressor-effect on richness was found in water, sediment, plant and animal microbiomes. Pairwise effects of time point and concentration were not significant due to lack of statistical power across all implied treatment levels. Similar results were obtained analysing Shannon diversity, except that Tb-mediated stress also increased diversity in the water, plant and animal microbiomes (**Figure S5B**). Overall, alpha diversity decreased in soil microbiomes by the three chemical stressors, while only Tb but not Bx and As affected Shannon diversity in most of the other food chain microbiomes.

**Figure 2.**
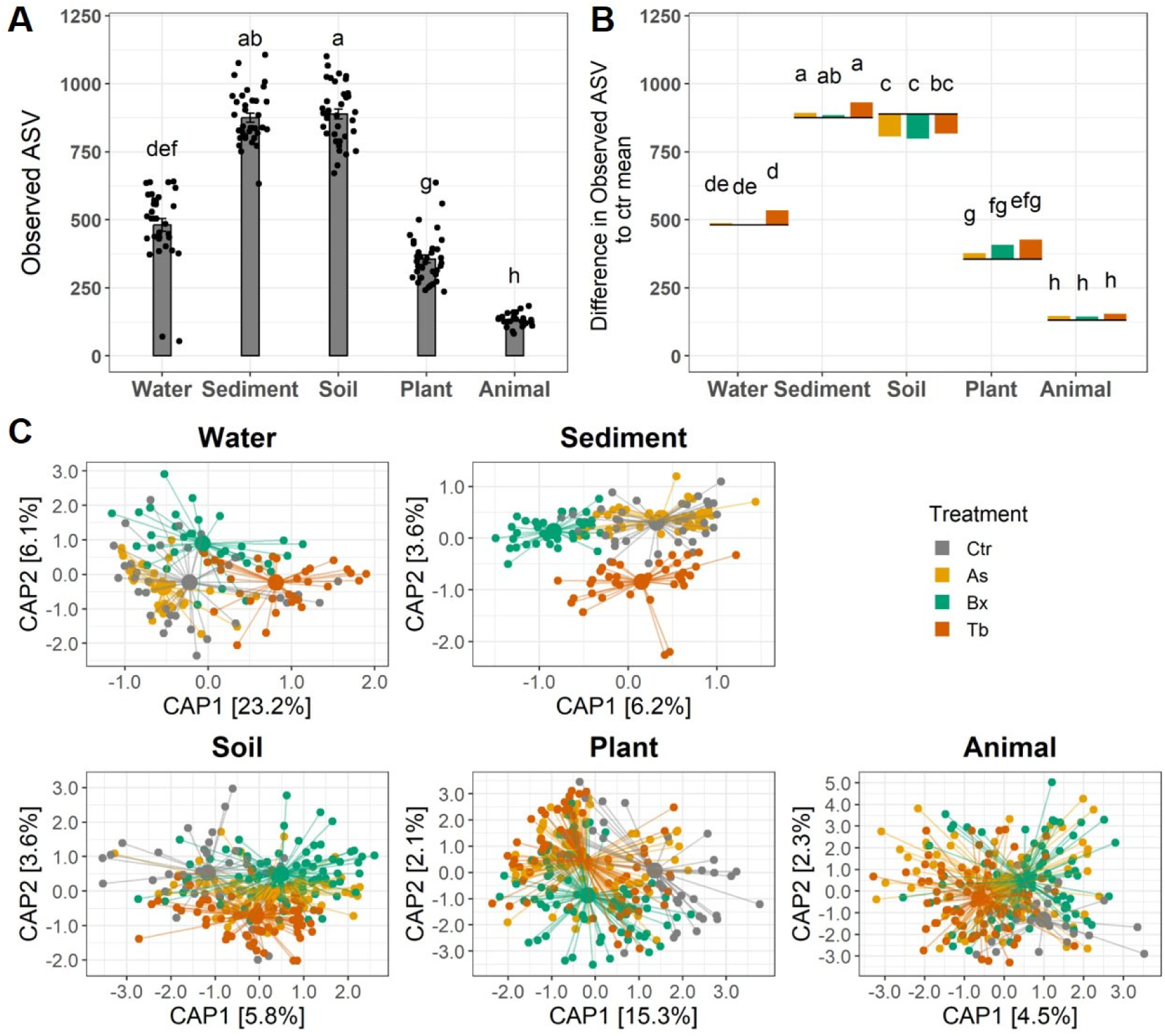
Observed ASV richness in control (**A**) and their relative changes in stress-induced treatments (**B**). In A) the jittered dots represent individual values of samples, while error bars indicate standard deviation per treatment. In B), differences to the means for each treatment (As, Bx, Tb) relative to the mean of their respective controls, which are represented as grey lines, are represented as barplots. The graphs are annotated with the Tukey HSD differences indicated by different letters (*P*<0.05). The same compact letter display is used as for the left panel. **C**) The first two significant axes of constrained ordination (dbRDA) are displayed for each component, with sample centroids per treatment indicated as different colours (see legend). Percentages of explained variance by each principal axis are indicated in square brackets.

### Chemical stressors alter microbiome composition to different levels

Next, we investigated whether and if to which magnitude the composition (beta diversity) of food chain microbiomes are perturbed by the chemical stressors. Besides the major differences between the food chain components (**Figure S4, Table S2**), much smaller, yet significant effects sizes were detected for time point, chemical treatment and concentration (all *P*=0.001). Similar to the alpha diversity analyses, most factor interactions were significant, but of very low effect sizes (∼1%). Post-hoc pairwise PERMANOVAs performed on the Bray-Curtis metrics between treatments revealed significant stressors-dependent decreases in beta diversity in water, soil and plant microbiomes (**Figure S6, Table S4**). Average beta diversity increased in animal microbiomes with all three stressors, while it increased after As and Tb treatments in sediment microbiomes but decreased after Bx treatment. We utilized constrained ordination of the Bray-Curtis metrics to visualize the chemical stressors-induced shifts in microbiome composition in comparison to the control groups (**Figure 2C**) and noted that performed models with treatment as an explanatory variable was significant (all *P<*0.01) in each component (**Table S5**). The ordinations further indicated that treatments altogether affected microbiome composition to different levels, with stressors impacts larger for the microbiomes in water (29.3% of explained variation; both axes) and plant (17.4%) than for those in sediment (9.8%), soil (9.4%), or animal (6.8%). The homogeneity of group dispersions tests showed significant dispersions (PERMDISP) in the low diversity microbiomes of plant (*P*=0.048) and animal (*P*=0.001), but not in the high diversity microbiomes of water (*P*=0.283), sediment (*P*=0.158), and soil (*P*=0.137) (**Table S6**). Overall, the three chemical stressors perturbed community composition of all tested food chain microbiomes. They generally caused the microbiomes to become more similar to each other, hence reduced beta diversity, with the exception of the animal microbiome, where the stress-perturbed microbiomes became more divergent than the control microbiomes. Furthermore, the low diversity microbiomes showed significant dispersion effects compared to high diversity microbial components of the food chain.

### Chemical stressors are associated with differential abundance of specific bacteria

We identified for each component the ASVs that differed significantly in mean relative abundance due to the chemical stressor treatments (i.e., contrasts between Ctr-As, Ctr-Bx, Ctr-Tb; *P*≤0.05) using negative binomial-based Wald tests (**Database S1**). Overall, the number of these unique, stressor-sensitive ASVs ranged from 1 (plant), 14 (water) 51 (soil), 65 (animal) up to 88 ASVs in the sediment microbiomes (**Figure 3A**). Consistent with the little overlap between the microbiomes (**Figure 1**), most stressor-sensitive ASVs were unique to a food chain microbiome, except one ASV that was common between water and sediment components. Within food chain components, 15/51 out of significant, unique ASVs in soil and 9/65 ASVs in animal microbiomes were commonly influenced by all three stressors while most other stressor-sensitive ASVs changed in abundance only after one and a few ASVs after two treatments. Stressor-sensitive ASVs rather decreased in abundance in water microbiomes, while in soil and animal microbiomes they mostly increased at the expense of few strongly decreasing ASVs (**Figures 3B**, **S7**).

**Figure 3.**
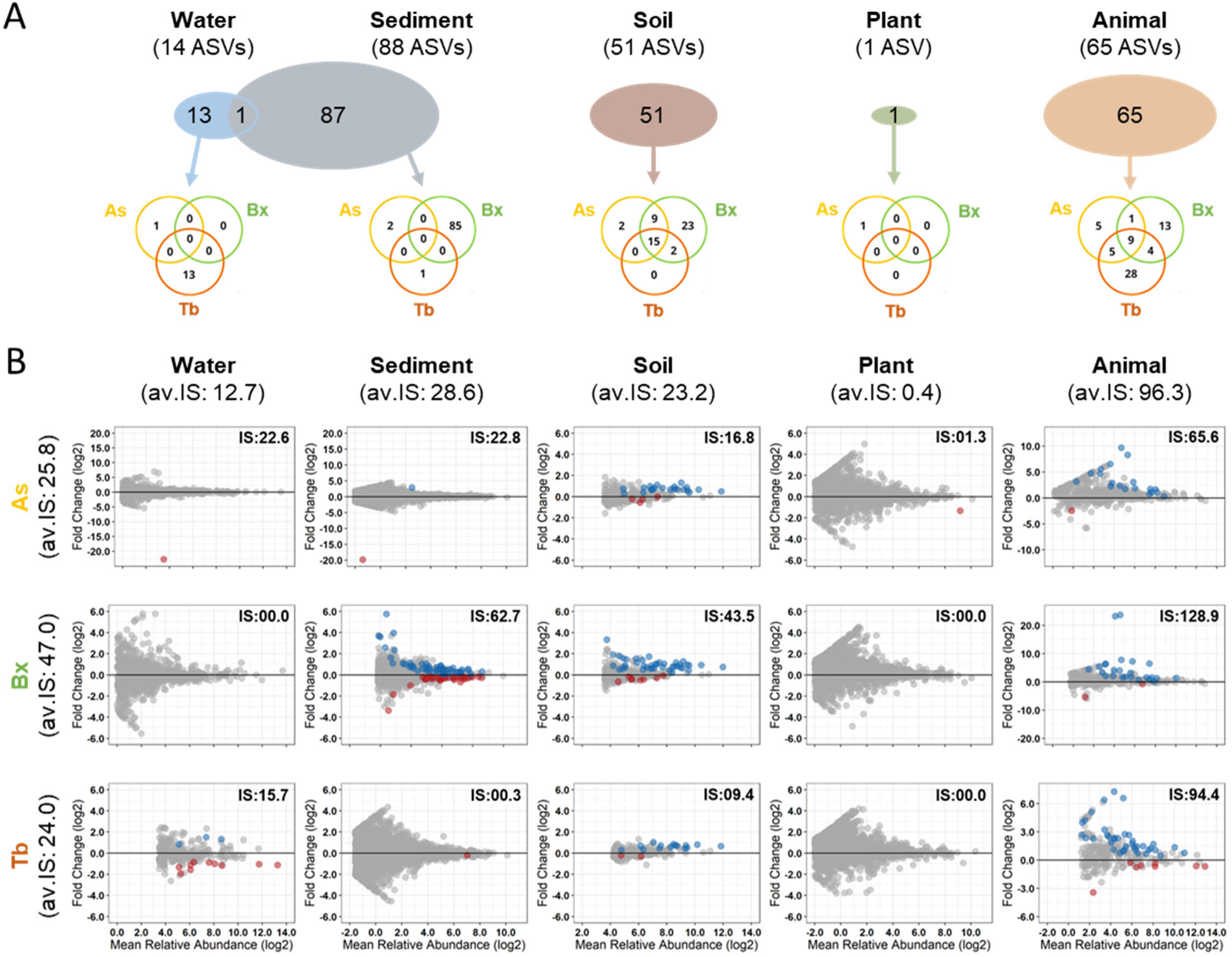
(**A**) Summaries of the stress-sensitive ASVs in the different microbiomes to the total number of stress-sensitive ASVs (circles scaled to their numbers) per component. The lower Venns detail the stress-sensitive ASVs by stress treatments As, Bx or Tb. (**B**) The MA plots display the log2-fold change of all ASVs and their log-mean abundance plotted on y- and x-axes, respectively, for each stress in each food chain component. ASVs being differentially abundant between control and treatments (As, Bx, Tb) were determined by DESeq2 analysis (Benjamini–Hochberg correction, *P*≤0.05). Colours refer to enriched ASVs in control (blue) or treatment (red) groups for all comparisons. Influence Score (IS) for each comparison was shown representing both the number of affected ASVs (*P*≤0.05) and their relative change in abundance for a given comparison (see method sections and Figure S7 for details). Average IS (av.IS) are indicated for each component.

We calculated an Influence Score (IS) to compare the impacts on differentially abundant ASVs across treatments and component. For a specific contrast (e.g., Ctr-As), the score considers both the number of affected ASVs and their log-fold change (see methods). Per food chain component, the highest average IS (av.IS) was noted for animal followed by sediment, soil, water and plant microbiomes (**Figure 3B**). Comparing the individual chemical stresses, the Bx treatment had higher average IS compared to As and Tb treatments. The IS were microbiome- and treatment-specific with the highest IS recorded for the Bx treatment on the animal microbiome, and lowest for the Tb and Bx treatments of the plant microbiome. Overall, the individual bacteria of the animal microbiome were generally most sensitive to the chemical stressors, in particular in the Bx treatment.

### Diverse bacterial taxa respond to the different chemical stressors

Next, we inspected the taxonomies of the stressor-sensitive ASVs in all food chain components. In the water microbiome, we found one and 13 ASVs specifically responding to As and Tb treatments, respectively (**Figures 3, S7**). Most of the ASVs that decreased by the Tb treatment belonged to Methylomonaceae (n = 3 ASVs) and Methylophilaceae (n = 4 ASVs) families. For sediment, we found two, 85 and one ASVs significantly changing with As, Bx and Tb treatments, respectively (**Figures 3, S7**). Of the Bx treatment, many shifting ASVs belonged to Syntrophaceae (n = 7 ASVs), Bacteroidetes (n = 6 ASVs), Anaerolineaceae (n = 5 ASVs) and Lentimicrobiaceae (n = 5 ASVs) families. For soil, we observed 26, 49, 17 ASVs significantly differing in abundance after As, Bx and Tb treatments, respectively (**Figures 3, S7**). In all three comparisons, most ASVs showed increase in abundance and few decreased. The increase was mainly ASVs from the Flavobacteriaceae family, specifically from *Flavobacter* genus (As, n = 7 ASVs; Bx, n = 18 ASVs; Tb, n = 7 ASVs), of which 6 ASVs commonly increased in all comparisons. ASVs from Burkholderiaceae (As, n = 4 ASVs; Bx, n = 5 ASVs; Tb, n = 2 ASVs) and Xanthomonadaceae (As, n = 3 ASVs; Bx, n = 4 ASVs; Tb, n = 1 ASV) also increased in all comparisons. One ASV from the Latescibacteria generally decreased in abundance in all treated groups. In the plant microbiome, only one abundant ASV belonging to the *Duganella* genus decreased in abundance upon As treatment, whereas no other ASV changed in abundance due to Bx and Tb treatments (**Figure 3**). Finally, in the animal microbiome, we observed 20, 27, 46 ASVs significantly differing in abundance after As, Bx and Tb treatments, respectively (**Figures 3, S7**). Most ASVs increased in relative abundance after treatment. This increase was associated mainly with ASVs from Lachnospiraceae (As, n = 11 ASVs; Bx, n = 13 ASVs; Tb, n = 18 ASVs), four of which were common in all treatment groups and belonged to *Lachnoclostridium*, *Shuttleworthia*, *Acetatifactor* and an Lachnospiraceae bacterium. ASVs from Ruminococcaceae (As, n = 3 ASVs; Bx, n = 4 ASVs; Tb, n = 12 ASVs) and Muribaculaceae (As, n = 2 ASVs; Bx, n = 4 ASVs; Tb, n = 6 ASVs) families also showed shifts in treatment groups. In general, the stressor-sensitive ASVs of the different microbiomes belonged to diverse taxonomic groups. In few cases, multiple ASVs of the same families had the same responses like Methylomonaceae and Methylophilaceae decreasing by the Tb treatment in water, or Flavobacteriaceae, Burkholderiaceae and Xanthomonadaceae increasing in soil in response to all 3 stresses or a consistent increase of Lachnospiraceae in the animal microbiome.

### Chemical stress disturbs bacterial co-occurrence networks

Co-occurrence networks were generated for each treatment and all microbiomes to investigate whether ASV co-occurrence may change due to a chemical stressor. In general, number of nodes and edges decreased from soil, sediment, water, plant to the lowest complexity network of the animal microbiome (**Figures 4A, S8**). The average edges per node were also the highest for soil, followed by sediment, water, plant, and the lowest for animal networks (**Table S7**). Positive, rather than negative associations were more prevalent in all networks (**Figure S8**). Several parameters of network complexity, such as node degree, betweenness centrality, closeness centrality, transitivity, were significantly different between control versus treatment groups (**Table S7**), and presented component-specific trends: For example, average node degrees decreased in most microbiomes after chemical treatments leading to sparse networks except for animal and sediment, where node degrees increased in the treated groups (**Figure 4B; Table S7**). We then examined the shapes, i.e. the distributions of the network’s node degrees and tested whether they were altered by the chemical treatments. From the evaluated different distribution types (power law, log normal, exponential and Poisson), the low complexity networks of animal and plant generally fitted best to a log-normal distribution while none of the tested data distribution types fitted to the high-complexity networks of water, sediment and soil (**Table S7**). However, no noticeable differences in degree distribution shapes were found after chemical treatments (Kolmogorov-Smirnov tests at *P*<0.05; **Figure S9**). Microbiome networks can also be used to detect hub nodes, which represent the most connected and possibly influential members of a given network. Based on Kleinberg’s hub centrality scores^54^, few hubs were observed in the lower complexity animal and plant networks, whereas higher numbers of hubs were observed the higher diversity components water, sediment and soil (**Figure 4, Table S7**). Chemical stressors consistently decreased the numbers of hub nodes for animal, plant and sediment components. In the soil microbiomes, however, the number of hub nodes increased in all treatment compared to control networks. For water, As treatment increased the number of hubs, while Bx and Tb treated microbiome showed lower numbers of hubs than the controls. Overall, chemical stress decreased network complexity for most microbiomes (water, sediment, soil and plant) except for the animal microbiomes where network complexity increased.

**Figure 4.**
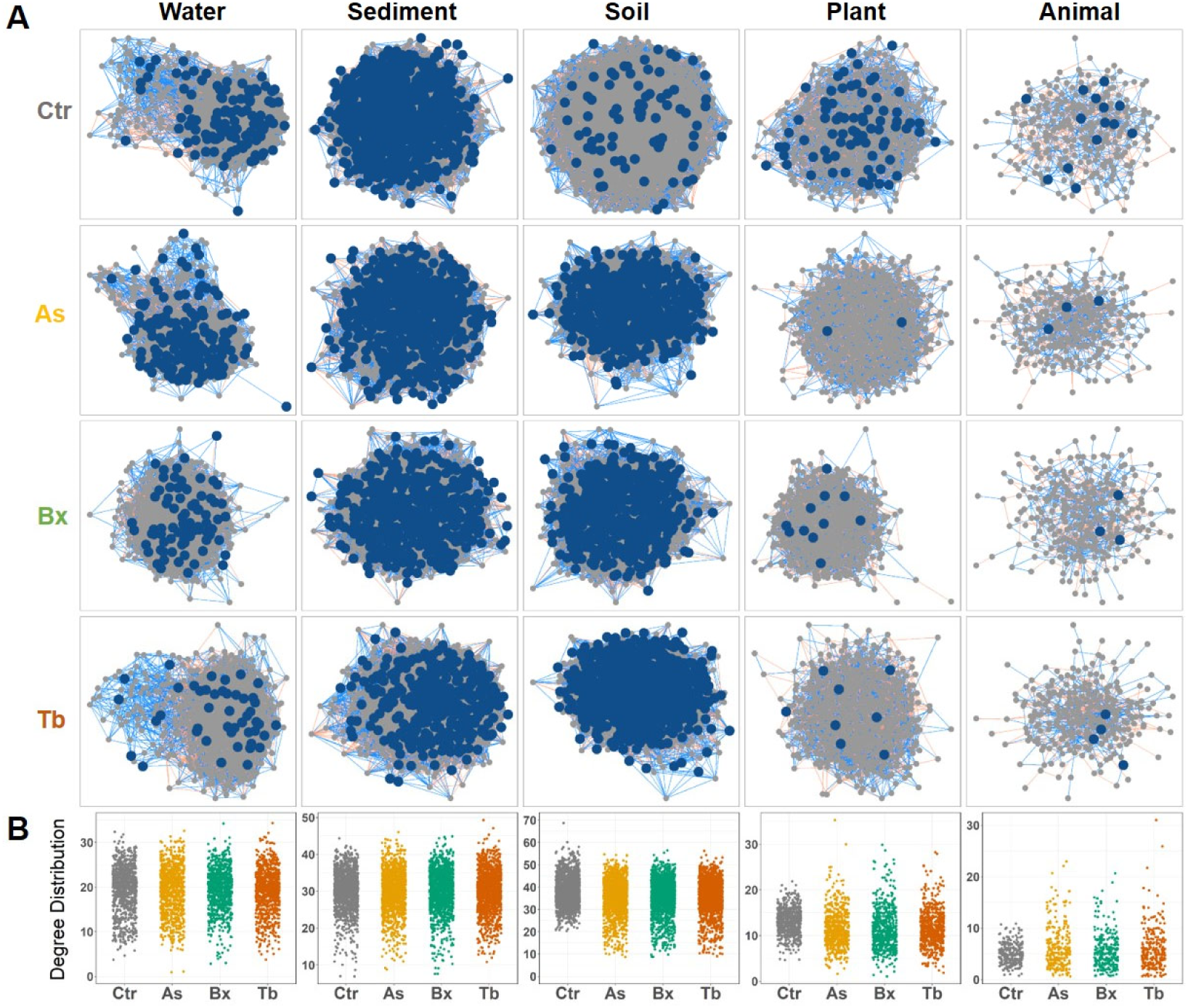
Co-occurrence network analysis. (**A**) Microbial association networks based on Meinshausen-Buhlmann method in SPIEC-EASI analysis for control versus treatments for all components. Nodes represent different ASVs, with blue colour nodes indicating hub nodes that are more connected to other nodes in the network (Kleinberg’s hub centrality scores >0.7). Blue edges indicate positive associations between ASVs, while red edges indicate negative associations. (**B**) The dot plots display the node degree’s (number of edges per node) of each ASV within the network as a function of treatment for each sample type. Differences in mean degrees for each chemical treatment vs. control were all significant (P≤0.001) in each component based on 10,000 bootstrap replicates of the underlying network properties (**Table S7**).

## Discussion

Microbes - whether mutualistic, commensal or pathogenic - have important roles in the health of a system as they are omnipresent with different communities in the different system components. A major gap towards a One Health understanding of microbiomes in a multi-component system is how sensitive or resistant different microbiomes are to different stresses. Are there commonalities and/or differences in the stress responses of different microbiomes to different stresses? To address this question, we systematically exposed different microbial communities of a multi-component system to three distinct chemical stressors at the same concentrations and we then analysed the microbiomes after the same exposure time. The system was an idealized food chain composed of water, sediment, soil, plants and animal microbiomes (**Figure 1A**). The three chemical stressors, i.e., the toxic trace element As, the bioactive plant metabolite Bx and the herbicide Tb were chosen because they can negatively impact the health of individual food chain components (As^13^, Bx^14,58^ and Tb^15,16^) and/or their microbiome^17–19^. Overall, we found that each component’s microbiome responded specifically to the different tested chemical stressors. Below we discuss various microbiome metrics to answer the main question of this study – commonalities and differences in stress responses of different microbiomes to different stresses – with the goal to identify the Achilles’ heel (i.e., the most stress-sensitive microbiome), as well as the most stress-resistant microbiome in our experimental food chain.

### No common stress responses of different microbiomes in their alpha diversity

We first discuss how the chemical stresses impacted the alpha diversity within and across the microbiomes of the experimental food chain. We confirm that free-living microbial communities (i.e., soil, sediment) have higher diversity and higher species richness than host-associated communities (i.e., plant roots, animal guts; **Figures 2A, S5A**), which has been shown earlier ^20,44^. The effects on alpha diversity by the three chemical stressors were not linked to whether communities have high or low levels of richness or diversity. The three applied chemical stressors reduced bacterial richness (**Figure 2B**) and Shannon diversity (**Figure S5B**) in soil but not in the other food chain microbiomes. This consistent decrease in soil bacterial alpha diversity by chemical stress is consistent with earlier work investigating the effects of individual chemical stressors on the soil microbiome^19,32,35,59,60^. Mechanistically, one could imagine that many or abundant bacteria, which tolerate and/or benefit from the chemical stressors, that they increase in abundance^26^ and that then leads to a decrease in overall diversity. The observed fold change in ASV abundances supports this idea (**Figure 3B**).

We further noticed that while As and Bx did not have any effects on alpha diversity, Tb-mediated stress increased Shannon diversity in the low diversity water, plant and animal microbiomes (**Figure S5B**). For such stress-specific changes, it could be postulated that some abundant taxa may be specifically susceptible to the compounds present in the chemical treatments and therefore, they decrease in abundance, what then allows other bacteria to proliferate increase overall diversity. Support for this postulation is seen in **Figure 3B**, where particularly the abundant bacteria were decreasing in abundance in the water and animal microbiomes. Finally, alpha diversity of the sediment microbiome remained fully unaffected (**Figures 2B, S5B**). One possible explanation that the chemical stressors did not affect these microbiomes could be that our study was limited to a duration of one week. One week may have been too short for slow metabolizing bacterial communities, such as those in sediments^61^, to result in detectable changes in alpha diversity. Regarding the main question of this study, i.e., how the chemical stresses compare in their impacts within and across the food chain microbiomes, we can conclude that stress effects on microbiome’s alpha diversity were both food chain component and chemical stress dependent. In other words, we did not find commonalities in the microbiomes’ stress responses from different food chain components.

### Food chain microbiomes responded deterministically to chemical stress responses and when host-associated combined with stochastic effects

Second, we reflect on the chemical stress impacts on community composition (i.e., beta diversity) that we found both, within and across the microbiomes of the experimental food chain. Consistent with alpha diversity, we found a strong “component” effect in beta diversity (**Figure S4**). This is expected as each component harbours compositionally different sets of bacteria and in different proportions^20,62^. In general, the three applied chemical stressors decreased average beta diversity of water, soil and plant microbiomes, while it increased in the mouse microbiomes (**Figure S6**). Minor changes were found in the sediment microbiomes, where it mildly increased or decreased according to treatment type. Individual studies of individual components (water^57^, sediments^63^, soils^19^, plants^35^ and animals^26^) may have suggested such heterogeneous changes in beta diversity in response to different stressors. Here, by comparing three stressors on five microbial communities, we demonstrate that the same stressors, in terms of chemical quality and quantity, have differential influence on different microbiomes. This systematic examination allows now to conclude whether stressors induced either deterministic, stochastic or a combination of these effects on microbiome composition. With deterministic effects, all microbiome members shift to new composition states without any dispersion effect (statistically: PERMANOVA and PERMDISP tests would be significant and nonsignificant, respectively). In contrast, with stochastic effects all microbiome members randomly disperse from their original composition state (PERMANOVA: nonsignificant; PERMDISP: significant). Third, there could be a combination of deterministic and stochastic effects where only some microbes move to a new community composition state, while others remain (PERMANOVA: significant; PERMDISP: significant). In conclusion, for the three chemical stressors we found deterministic changes in water, sediment and soil microbiomes and in plant and animal microbiomes, the detected deterministic changes were combined with stochastic effects in dispersion (**Tables S4, S5**).

A caveat for this conclusion is that deterministic and stochastic effects can vary with time and stress strengths: For instance, mild stress can lead to an increase, but severe stress leads to a drastic reduction in beta diversity compared to that of healthy subjects^64^, as also shown here for most microbiomes of the experimental food chain. However, we could not evaluate the effects of chemical concentrations as well as of time point due to a statistical limitation (PERMDISP does not allow interaction terms) and due to lack of statistical power to resolve the significance of each pair of combined treatment levels. Our experiment was designed to systematically compare all components along the food chain with different chemical stressors and allowed to highlight that any stressor effect on a component microbiome could not be generalized to other microbiomes of the food chain. Future work is needed to reveal fine-grained differences of combinations of chemical concentrations and temporal changes. Also host effects on corn roots or mouse guts should be accounted in such interactions. This research would aim to understand how the microbiome evolves over time, especially in terms of the resilience and resistance of microbial communities following initial dysbiosis induced by different chemical concentrations.

### Stress-specific microbiome changes may result in health effects

Because the microbiomes of our experimental food chain do not share much overlap in bacterial species (**Figure 1B, 1C**), it is of little use to discuss taxonomic commonalities and disparities of the microbiome members that responded to the different stressors. Instead, we explored whether the taxonomic information of the stress-sensitive ASVs in a given microbiome, may be indicative for eventual health effects on the food chain component. For this we focused on the major stress-sensitive ASVs in each microbiome. Only one ASV in plant and few stressor-sensitive ASVs in the water microbiomes were detected, while several stressor-responsive ones were found in sediment, soil and animal microbiomes (**Figure 3A**). With the exception of a single ASV after As stress, no changes were observed in the corn root microbiomes after the stress treatments. Albeit negative health effects had been described for plants^25^, this finding may indicate that the root microbiome may be relatively insensitive or slow to stress perturbation compared to the other components. In the water microbiome, ASVs of the Methylomonaceae and Methylophilaceae mainly decreased in abundance (**Figures 3, S7**). Members of this family are responsible for methane oxidation in lakes and are important members of lake microbiome. Thus, their decrease in after Tb treatment could indicate a disruption of normal methane cycling in the water microbiome^65^ and may point to a negative health effect. The major effect on the sediment microbiome was observed in response to the Bx treatments with several shifting ASVs belonging to the Syntrophaceae, Bacteroidetes, Anaerolineaceae and Lentimicrobiaceae (**Figures 3, S7**). Members of these families are abundant in sediments and are often associated with bioremediation, organic matter decomposition and acetate oxidation processes^66–69^. However, future experiments are needed to test if their change in abundance affects sediment health.

For the soil microbiome, the majority of ASVs responded with an increase in relative abundance, particularly after As and Bx treatments and several of these ASVs were members of Flavobacteriaceae, Burkholderiaceae and Xanthomonadaceae families (**Figures 3, S7**). Members of the Flavobacteriaceae are dominant in soil and marine microbiomes, but are also found in association with plant roots. Specifically, the genus *Flavobacter* is specialized in uptake and decomposition of organic matter due to its capacity to hydrolyse organic polymers^70,71^ and therefore, their wide biotechnological use in biotransformation, wastewater treatment and bioremediations^71^. Similarly, members of Burkholderiaceae, specifically the *Massilia* genus can degrade herbicides, metabolize aromatic hydrocarbons and are resistant to metals^72–74^, thus their increase in relative abundance after the stress treatment. Members of the Xanthomonadaceae, mainly *Lysobacter* bacteria possess antimicrobial and antifungal properties, secret many bioactive compounds, are resistant to arsenite, and function in bioremediation of hydrocarbon polluted soils^75^. Similar as for sediments, the shifts of the bacteria in response to the chemical treatments, are consistent with metabolic traits, but whether their change in abundance in the microbiome affects soil health remains to be experimentally assessed.

The major effect observed in the animal microbiomes was that ASVs from Lachnospiraceae, Ruminococcaceae and Muribaculaceae increased after the stressor treatments (**Figures 3, S7**). Lachnospiraceae and Ruminococcaceae are two commensal families specialized in the degradation of complex plant material, but they may also provide protection against enteric infections in the human gut. Some Ruminococcaceae and Lachnospiraceae are butyrate producers, an important source of energy for gut epithelial cells, and they support humans to maintain epithelial barrier integrity and thereby, prevent diarrhea^76,77^. Increase of both of these families after exposure of humans to toxic trace elements and their beneficial roles in the gut health was found earlier^78,79^. The Muribaculaceae family commonly occurs in animals with high abundance in rodents and provide several important functions to the host^80^. Interestingly, members of Muribaculaceae were found to be associated with enhanced longevity in mouse^81^. Hence, the taxonomic information of the stress-sensitive ASVs clearly point to health effects on the animal host.

Taken together, although some of the stressor-specific influences on the different microbiomes indicate individual health effects, a next step is now to compare systematically the health effects, both within and across the components of the experimental food chain.

### Chemically stressed microbiomes become structurally sparser

Finally, addressing the main question of this study – commonalities and differences in microbiome’s responses to different stresses - we specifically investigated the stress-induced changes in network properties, as these can reveal hidden patterns in the communities usually not captured by diversity metrics^82^. Generally, the inferred networks reflected microbial diversity with the number of nodes and edges among microbiome members of a given food chain component. As expected from bacterial richness and diversity, number of nodes and edges decreased from soil, sediment, water, plant to the lowest complexity network of the animal microbiome (**Figures 4A, S8**). We found that positive associations outnumbered negative associations when analysing networks from five different components (**Figure S8**). This did not change with chemical stress, which suggests that microbial community changes are primarily driven by conjointly enhancing biological fitness rather than by increasing competitive pressure. Chemical stress still changed several network parameters including the distribution of node degrees (**Table S6**). Node degree suggests how well a node is connected to other nodes, its decrease suggests loss in bacterial community cohesiveness and overall sparser network structure and instability. With the exception of the mouse microbiome, we observed that all networks became structurally sparser after applying chemical stressors (**Figure 4**). In addition, we noticed changes in abundance of hub nodes also called keystone taxa, which showed high degree connectivity to other nodes and are considered as important members of the community^83^. After chemical stress, the number of keystone taxa decreased in most components’ networks, except in soil where they increased. Such a decrease suggests losing contributions of important taxa, which can potentially decrease the community stability and affect health of the overall community. Such decrease in keystone taxa in response to chemical stress is in accord with previous studies investigating chemical fertilizers or pesticides^84^. Overall, our performed co-occurrence analysis revealed that network properties changed after the chemical treatments in all components and with all stresses. Networks became often sparse with loss of keystone taxa, which could negatively influence the resilience of each component and indicate dysbiosis.

### Conclusions

The main motivation for this study was to answer whether different microbiomes cope with different stresses with common and/or differential stress responses. We can conclude from applying three representative chemical stressors to five microbiomes over a short time typically found along a human food chain, that each microbiome responded in its own way to stress treatments. We found stress and microbiome-specific shifts in community composition with some of the changing members pointing to possible impacts on food chain health. The shifts to different dysbiotic microbiomes, that we observed, are reminiscent of the Anna Karenina principle^9^. It refers to Leo Tolstoy’s dictum that “all happy families are alike; each unhappy family is unhappy in its own way” and applied to microbiomes, it states that dysbiotic individuals vary more in community composition than healthy individuals. In addition to specific responses on diversity and community composition, our work revealed that chemical stress commonly affected the complexity of bacterial co-occurrence. Most microbiome networks became sparser with fewer keystone taxa, while stress increased these properties in soil networks. Hence, chemical stressors induce microbiome alterations that may differentially impact the stability and structure of the different microbiomes along a food chain. A goal of this study was to identify the Achilles’ heel of our experimental food chain. With reference to the influence score, which takes number and abundance changes of ASVs into account, the animal gut presented the most stress-sensitive microbiome in our experimental food. However, extending the Anna Karenina principle to the wider One Health context, implies that each component’s microbiome will have its own Achilles’ heel and therefore, investigations that particularly elucidate the contribution of microbiomes to the health of a system are needed.

## Supporting information

RDS R object (see Supplementary Material)

## Acknowledgments

We thank Evelyne Vonwyl for her help in the laboratory and field work and Luyao Tu and Tobias Schneider for their help in the field as well as Anna Muntwyler (all from the Department of Geography, University of Bern). For experimental and logistical support, we are also grateful to Oliver Schären, Luca Beldi, Disha Tandon, Stephen Leib, Stefan Neuenschwander, and Miguel Terrazos Miani (all from the Institute of Infectious Diseases, University of Bern). We also thank Pascal Wyss, Christina Widmer, Bernardus CJ Schimmel and Lei Wang for support during sample collection (all from the Institute of Plant Sciences, University of Bern). The sequencing data were generated in collaboration with the Genetic Diversity Centre (GDC), ETH Zurich.

## Author contributions

The study was conventionalized and supervised by W, FR, SGV, MB, MG, SS, MA, AM, SH, ME, KS and AR. Funding was acquired and resources provided by FR, SGV, MB, MG, SS, MA, AM, SH, ME, KS and AR. Experiments were performed by W, ACH, CT, LT, VC, PM, MN, MM, TCC, FR and SGV. Specific methodology was developed and provided by W, ACH, MCC, PM, TCC, AM, ME, KS and AR. W, CT, KS and AR were responsible for data curation. Software was developed, data was analysed, validated, visualized and the first draft written by W, KS and AR. All co-authors have contributed to reviewing and editing the manuscript and agree on the final version of this study.

