## Supplementary text, figures and tables for "Component specific responses of the microbiomes to common chemical stressors in the human food chain"

Wasimuddin et al. (Year), Journal

#### Table of Contents

Supplementary Methods ... 2
- Water/sediment experiment ... 2
- Soil/plant experiment ... 3
- Animal experiment ... 5
- References ... 6

Supplementary Figures ... 7
- Figure S1 | Experimental setups ... 7
- Figure S2 | Validation of treatment solutions ... 8
- Figure S3 | Soil/plant experiment ... 9
- Figure S4 | NMDS analysis ... 10
- Figure S5 | Alpha diversity, Shannon index ... 11
- Figure S6 | Beta diversity ... 12
- Figure S7 | Stress-sensitive ASVs and Influence Scores ... 16
- Figure S8 | Network characteristics ... 17
- Figure S9 | Network degree distributions ... 18
- Supplementary Tables ... 19
- Table S1 | Basic parameters of the experimental soil ... 19
- Table S2 | Factors affecting beta diversity ... 20
- Table S3 | Factors affecting alpha diversity ... 21
- Table S4 | Pairwise factors affecting beta diversity ... 22
- Table S5 | Constrained ordination for treatment effect in each component ... 22
- Table S6 | Testing homogeneity of group dispersion ... 22
- Table S7 | Network properties ... 23

Supplementary Data ... 24

#### Supplementary Methods

##### Water/sediment experiment

*Source of lake water and sediment:* In March 2019, surface sediment (top 10 cm) was retrieved from the deepest (~22 m) part of Lake Moossee (47.0238°N, 7.4775°E), located 8 km north of the city of Bern, Switzerland, using an Ekman grabber. Simultaneously, 23 L of lake water were collected from the bottom of the lake using a battery-powered water pump with a depth-marked tubing. Sediment and lake water were transferred to the laboratory under cold and dark conditions and immediately stored in dark at 4°C until sample preparation.

*Setup:* Incubation vessels were prepared using 250 mL amber glass jars (I-Chem ™, Thermo Scientific, Switzerland) previously cleaned and backed (275°C) for 24 h to remove any presence of organic material. Incubation vessels designated for As treatments were first cleaned by acidification (10% $HNO_3$ for 24 h), rinsed thoroughly with ultrapure water (Milli-Q, Millipore) and dried over the hood.
The water and sediment samples were manually homogenized and filled in the incubation vessels containing 50 g of sediment and 160 mL of lake water in all vessels. Immediately after preparation, the incubation vessels were closed to keep the experiment under anaerobic conditions. At least 0.5 cm headspace was left at the top of each vessel to avoid sorption of tested substance to the cover. Incubation vessels were transferred to a control chamber (HPP 260, Memmert, Germany), and kept under dark conditions at 6°C for 7 days for acclimatization before adding the test substances. A total of 144 incubation vessels were prepared, consisting of 4 concentrations (0, 10, 100 and 1,000 µg/L) × 3 chemicals (As, Bx, Tb) × 2 time points (d1 and d7) × 6 replicates.

*Chemical application procedure:* A constant amount of 533 µL of the 300x-treatment solutions (3, 30, 300 mg/L, see above) was added to 160 mL of lake water and mixed with a glass stirring rod (Huberlab, Switzerland) without disturbing the sediment to reach final (1x) concentrations of 10, 100 and 1,000 µg/L. Similarly, the treatment solutions of Tb, but diluted in ethanol, were used and control samples adjusted to contain the same amounts of ethanol. We applied the chemicals in daily intervals to approximate continuous exposure of the microbiomes. Manipulations were conducted under anaerobic conditions using a glove box (Systemtechnik, Germany) to avoid disturbance to the incubation system and the vessels were immediately returned to the incubator after treatment. We confirmed the final concentrations of the three chemicals in the water of the water/sediment experiments by collecting samples one day after the first application (**Figure S2)**.

*Sampling:* The first sub-group of 72 vessels was harvested at the first time point d1 (one treatment application). The second sub-group of 72 vessels was harvested at d7 (second time point). At harvest, the water and sediment fractions were transferred to individual 50 mL polypropylene centrifuge tubes (Sigma-Aldrich, Germany) and stored at -20°C until analysis.

**Soil/plant experiment**

*Source:* The soil for this experiment was taken from "Q-Matte" in Frauenkappelen, Switzerland (46.9555N, 7.3324E) as described earlier[1]. Our analyses revealed that it contained simultaneous low baseline levels of As, Bx and pesticide residues (specifically glyphosate and Tb; **Table S1**). The upper A-horizon (20 cm) of agricultural soil was collected and stored as a large batch (24 $m^3$) next to our research facility in Ostermundigen for repeated experimental accessibility. Natural storage conditions were chosen by keeping the soil outdoors and seeded with a grass cover. Upon experimental needs, we could collect soil for the same experimental soil type[1]. Soil for this experiment was collected on the 4th of February 2019, sieved to 5 mm and mixed with 20% (v/v) autoclaved sand ("CAPITO" 1-4 mm, 25 kg, LANDI, Switzerland). Sand was added to facilitate water flow and enhance draining of the soil pots.

*Setup:* The experiment was conducted in custom-made pots of ca. 200 mL (12 cm height, 4 cm diameter). The pots consisted of open polypropylene tubes (cut from HT-tubes; Marley, Switzerland) and closed at the bottom with a thin gardener's net (Windhager Trennflies, Germany) using tape (TESA Professional Strong Duct Tape, Switzerland). This system was developed as it allows to control soil water content by placing the pots on household paper for drainage (see below; **Figure S3A**). The pots were filled with 150 g of soil:sand mix and watered with ~30 mL of tap water to reach 60% soil water content.
The corn (*Zea mays* L.) inbred line W22 (wild type) and the mutant *bx1* (in background W22[2]) were used in this experiment. Seeds were surface sterilized using commercial bleach containing 5% active hypochlorite (Potz Javel-Wasser Natur, Switzerland) for 6 min and rinsed with sterile distilled water five times. For germination, seeds were soaked in sterile distilled water for 8 h in the dark before transferring them on wet filter paper where they germinated overnight in the dark. One germinating seedling was planted per pot. Pots were randomly placed on a greenhouse table (light/dark 14/10 h, daily temperature from 14°C to 22°C, night temperature from 10°C to 14°C, humidity range 40-60%, 50,000 lx illumination). Plants were watered on a weight basis to keep water content between 40-60% and pots were randomized throughout the experiment. To avoid cross-contamination by eventual leaking treatment

solutions, lids of petri dishes were placed below the racks containing the pots trays so that they do not touch the bottom of the tray (**Figure S3B**). Plants were 10 days old (after germination) at the start of the experiment. A total of 240 pots were prepared and consisted of 10 treatment groups (Ctr at 0 µg/L; As, Bx and Tb, each 3 concentrations) × 2 time points (d1, d7) × 12 replicates (6 for W22, 6 for *bx1*). Of note, although this factor of plant genotype was part of the setup it was not examined in this study because it is nested only within the plant component and because we focused here on component-level differences across the experimental food chain.

*Chemical application procedure:* We applied the chemicals as 3x-treatment solutions (30, 300 and 3,000 µg/L, see above) to the pots because the soil already contained water, which limited the volume we could apply. To reach the final (1x) concentrations of chemicals as in the other systems (10, 100 and 1,000 µg/L), we specifically "cycled" the soil water content (SWC) between 40% and 60%, where the daily applied volume of the treatments corresponded to 20% of SWC (**Figure S1**). The SWC was controlled on a weight basis. Before treatment application, the SWC was reduced to 40% by draining the pots on household paper, and after treatment it reached 60% SWC again. This cycling of SWC in daily intervals allowed (i) repeated treatment application to approximate continuous exposure of the microbiota to the chemicals and (ii) such an "as-large-as-possible" volume permitted a more homogenous distribution of the compounds in the soil matrix. We confirmed the 3x concentrations as used for the soil/plant experiment by collecting samples from the treatment solutions (**Figure S2**).

*Sampling:* Two series of 120 pots were harvested at the two time points (d1, d7). The entire soil core including the root system was removed from the pots by inverting them, gently pulling the plant out and placing it on sterile square petri dishes (120 x 120 x 17 mm; Greiner, Germany) for fractionation. The soil core was cut to the -2 to -7 cm soil section with a sterile scalpel, from which soil and root parts were collected (**Figure S3C**). The 5-cm-long root fragments were taken with forceps into clean 50 mL plastic centrifuge tubes (Cellstar tubes, PP, blue screw caps graduated, conical bottom; Greiner Bio-One, Switzerland) for washing off the adhering rhizosphere soil (two washes with 30 mL sterile water and inverting the tubes 10 times), washed root fragments were gently dried by wrapping them between tissue paper sheets, transferred to 15 mL centrifuge tubes and snap frozen in liquid nitrogen until further analysis. The soil corresponding to the 5-cm-section of the soil core that remained on the petri plate was homogenized with a sterile spatula, and a about 1 g aliquot was collected in 2 mL labelled screw cap tubes (Sarstedt, Germany). Tubes were snap frozen in liquid nitrogen and stored at -80°C until further analysis.

**Animal experiment**

*Source:* Animal experiments were performed in accordance with animal experiment license BE44/18 approved by the Bernese Cantonal Ethical committee for animal experiments and carried out in accordance with Swiss Federal law for animal experimentation. Eight-week-old specific pathogen-free (SPF) C57BL/6J females were purchased and supplied from the same hygiene barrier (Charles River, Germany).

*Setup:* Animals were randomized at different stages at Charles River and once again on arrival at the experimental facility. Mice were kept at the rodent experimental facility of the Institute for Infectious Diseases, University of Bern, in sterile individually ventilated cages (five individuals per cage) under positive pressure (Tecniplast, Italy) and controlled ambient temperature (25–27°C) and relative humidity (52–60%). These conditions were maintained over the entire duration of the experiment. All mice were supplied with purified diet D11112201ii (Research Diets, USA) *ad libitum* upon arrival and kept on the same diet for the entire duration of the experiment. The D11112201ii diet had been sterilized by gamma irradiation (2x 10-20 kGy at Research Diets, USA), followed by 30-60 kGy x-ray irradiation (Steris, Switzerland), and sterility was confirmed by absence of cultivable aerobic and anaerobic microbial culture in faeces and cecal contents of sentinel germ-free mice that had been fed with this diet. Mice were provided with sterile drinking water *ad libitum* (autoclaved surgical irrigation water; Baxter, USA). A total of 100 mice were utilized for 10 treatment groups (Ctr at 0 µg/L; As, Bx and Tb, each 3 concentrations) × 10 replicates.

*Chemical application procedure:* After three weeks of accustoming the mice to the conditions, the animals were randomized to the ten treatment groups and the treatments with chemicals started. The 1x treatment solutions (10, 100 and 1,000 µg/L, see above) of the three chemicals were provided as drinking water. The treated drinking waters were sterile filtered (0.2 µm exclusion size) and provided *ad libitum* for 7 days to ensure continuous exposure of the mice to the chemicals. We confirmed the 1x concentrations by collecting samples from the drinking water used in this experiment (**Figure S2**).

*Sampling:* Fresh faecal samples were collected before treatment (time point d0) and after treatments at time points d1 and d7 from manually restrained animals and were snap frozen in liquid nitrogen and stored at -80ºC until further analysis. All animal handling was carried out under sterile conditions using surgical equipment under a sterile laminar flow.

#### Supplementary Figures

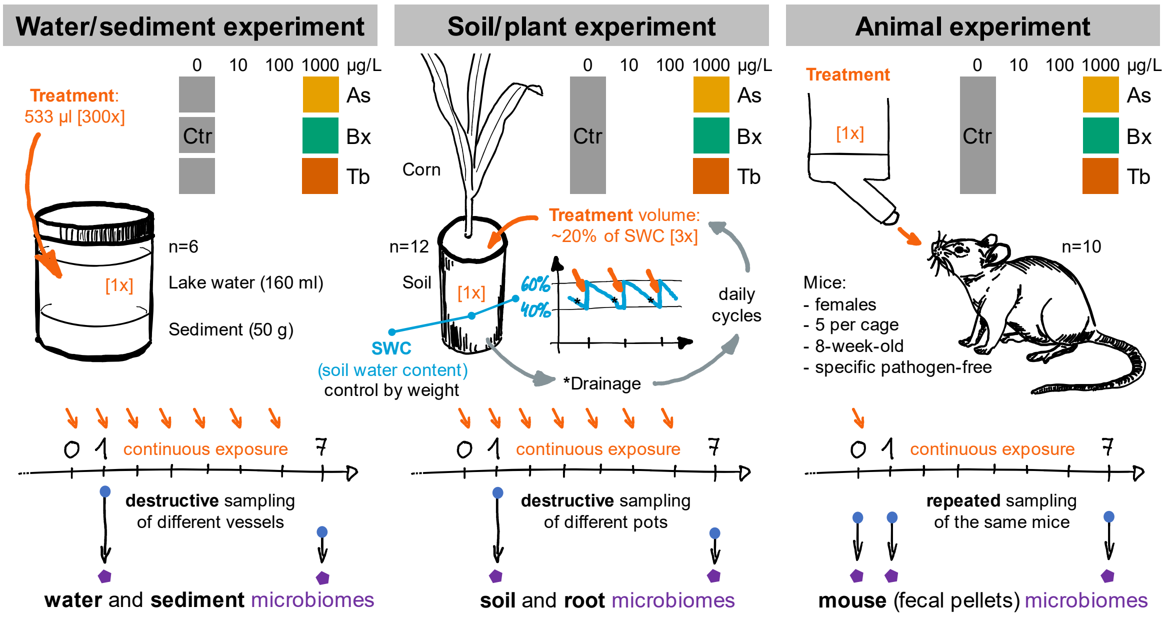


**Figure S1 | Experimental setups**
The model food chain consisted of water, sediment, soil, plant and animal components, which were treated with As, Bx and Tb stress in three parallelly executed experiments. These experiments were named the ‘water/sediment’, the ‘soil/plant’ and the ‘animal’ experiments. The chemical stress solutions, indicated in orange color, were applied from concentrated ‘treatment solutions’ to final concentrations of 10, 100 and 1000 µg/L and in daily intervals to reach as best as possible continuous exposure in ‘water/sediment’ and ‘soil/plant’ experiments. The chemical stress solutions were applied as final concentrations in the ‘animal’ experiment. The cartoons of the incubation vessels, corn growth system and the mice provide experimental details on relative proportions (water to sediment), the daily cycling of soil water content to reach 1x exposure, and specific growth setups and replicate numbers. The time lines at the bottom depict the destructive or repeated samplings of microbiomes at zero, one or 7 days after exposure.

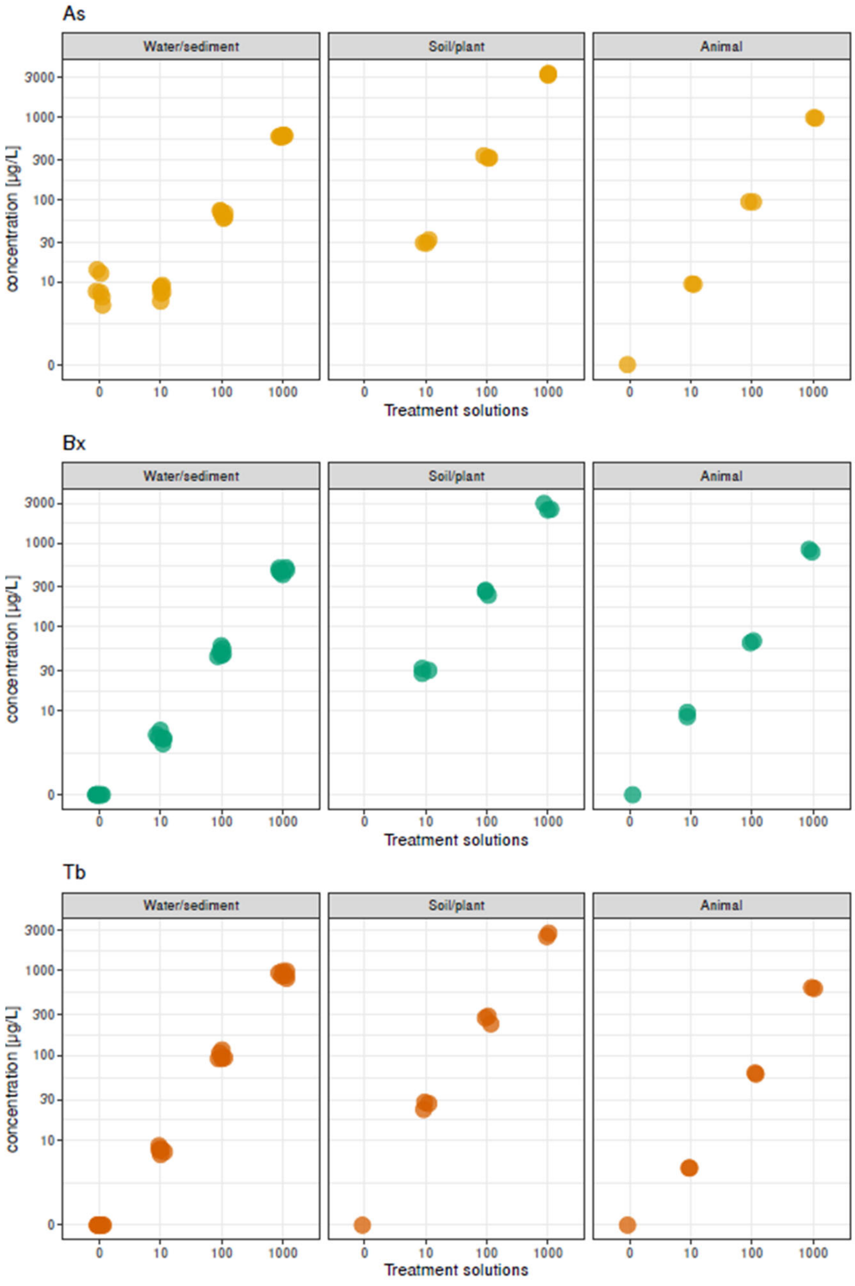


**Figure S2 | Validation of treatment solutions**
The actual concentrations of the As, Bx and Tb water in the water/sediment and animal experiments and 3x treatment solutions used in the soil/plant experiment were analyzed for the effective concentration of the chemical compounds. The control (0) and 10, 100 and 1000 µg/L waters and treatment solutions contained the standardized 10-fold increments of stress chemicals. Replicate numbers were n=6 for the water/sediment, n=2 to 3 for the soil/plant and n=2 for the animal experiments.

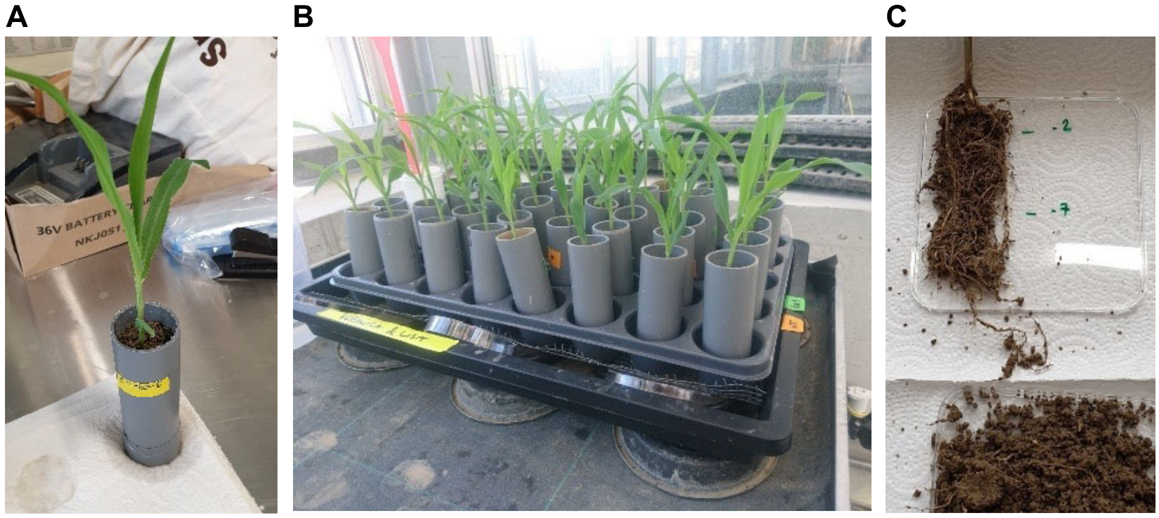


**Figure S3 | Soil/plant experiment**
**A**) The grey polypropylene pots were closed at the bottom with a thin gardener's net fixed with tape. The permeable bottom of the system allowed to control soil water content by placing the pots on household paper for drainage. Before application of the chemicals, we reduced the soil water content (SWC) down to 40% based on pot weight. With the application of the chemicals SWC reached again 60%. **B**) Lids of petri dishes were used to lift the racks containing the pots from the trays so that they do not touch the bottom of the tray. This was done to prevent eventual cross-contaminations by leaking treatment solutions. **C**) Sampling of the root systems on sterile square petri dishes where the root segment corresponding to the -2 to -7 cm soil section was cut with a sterile scalpel for microbiome analyses.

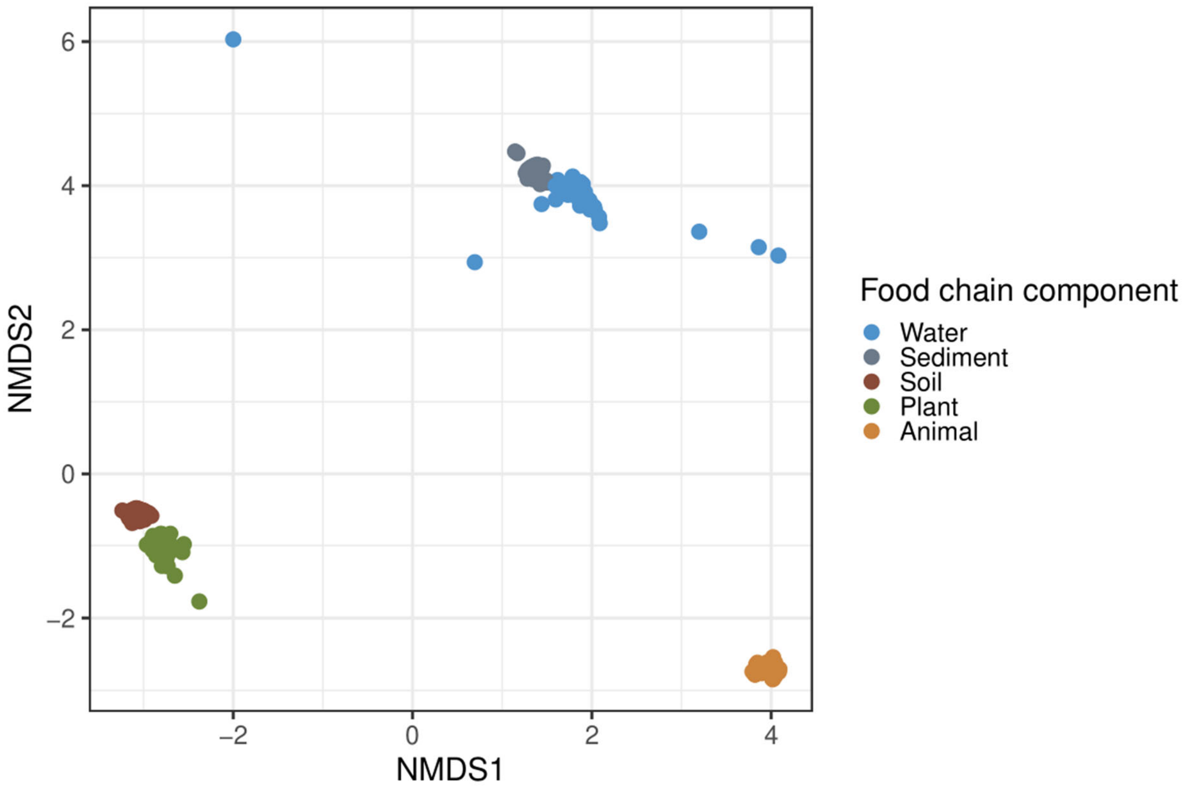


**Figure S4 | NMDS analysis**
Unconstrained ordination using non-metric multidimensional scaling (NMDS) based on Bray-Curtis dissimilarity indices. The NMDS analysis visualizes patterns of separation among the different sample groups (indicated by colors).

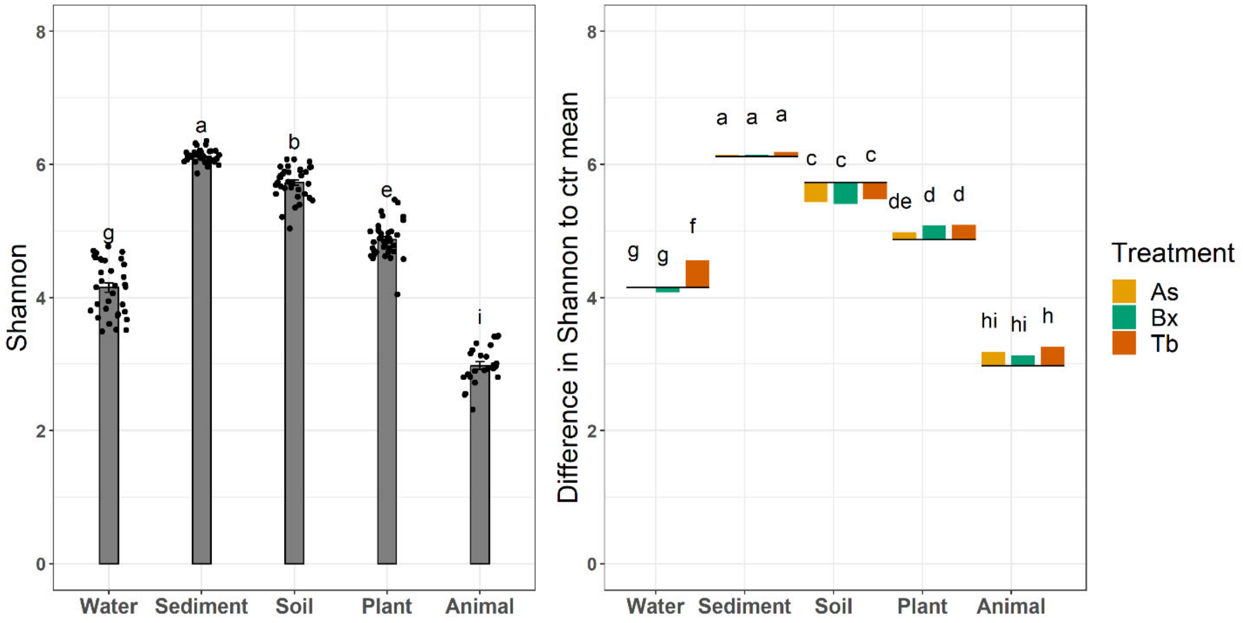


**Figure S5 | Alpha diversity, Shannon index**
(**A**) Alpha or within-sample diversity based on the Shannon index of the control-treated microbiomes. Jittered dots represent values of individual samples and error bars indicate standard deviation per treatment. **B**) The bars represent the differences of the means for each treatment (As, Bx, Tb, different colors) relative to the mean of their respective controls (represented as grey lines). The graphs are annotated with the Tukey HSD differences indicated by different letters ($P$<0.05). The same compact letter display is used as for A and B.

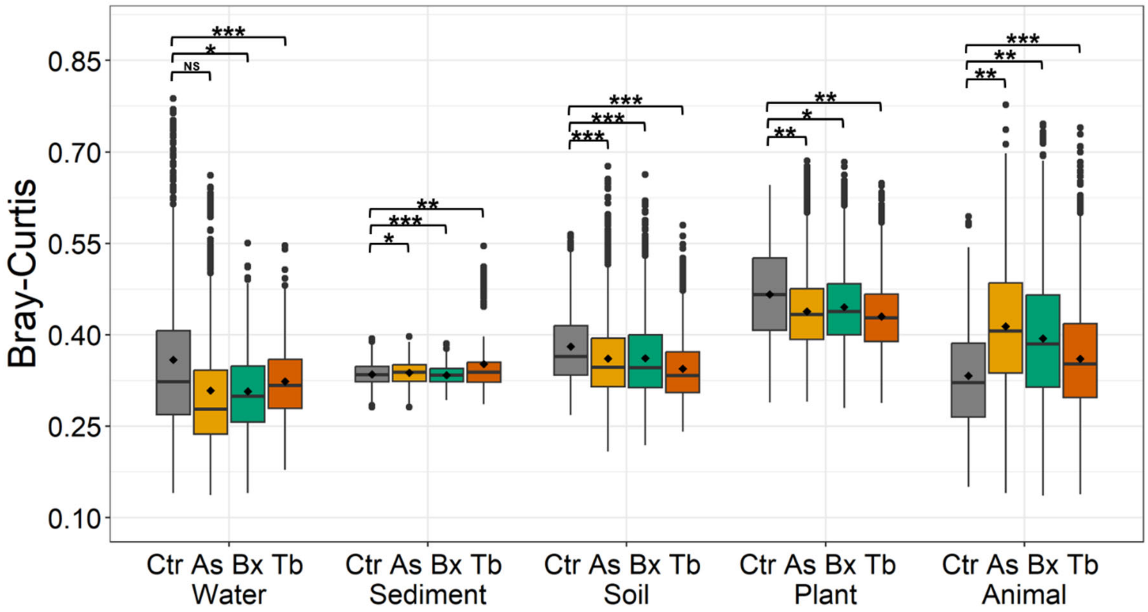


**Figure S6 | Beta diversity**

The boxplots display all Bray-Curtis distances within sample groups of control, As, Bx and Tb treated microbiomes for each component of the model food chain. Multivariate homogeneity of group dispersions (variances) was analysed by the function *betadisper* (R package *vegan*). To test if one or more treated groups is more variable than the control group, ANOVA of the distances to group centroids was performed using a permutation test (999 permutations), which permutes model residuals to generate a permutation distribution of *F* under the null hypothesis of no difference in dispersion between groups. Levels of significance include * for *P*≤0.05, ** for *P*≤0.01 and *** for *P*≤0.001.

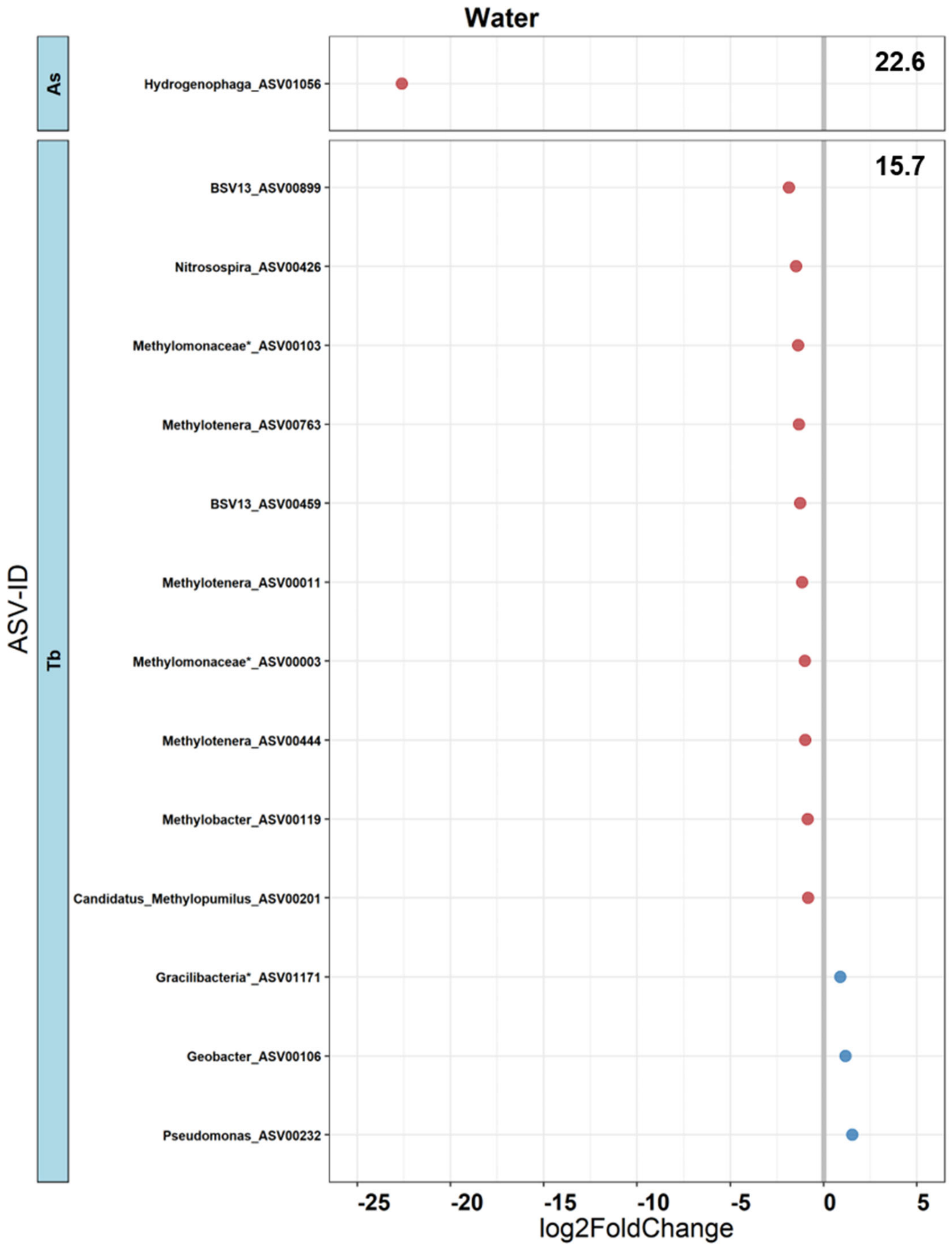
Water
As
Hydrogenophaga_ASV01056
22.6
Tb
BSV13_ASV00899
Nitrosospira_ASV00426
Methylomonaceae*_ASV00103
Methylotenera_ASV00763
BSV13_ASV00459
Methylotenera_ASV00011
Methylomonaceae*_ASV00003
Methylotenera_ASV00444
Methylobacter_ASV00119
Candidatus_Methylopumilus_ASV00201
Gracilibacteria*_ASV01171
Geobacter_ASV00106
Pseudomonas_ASV00232
15.7
ASV-ID
-25
-20
-15
-10
-5
0
5
log2FoldChange

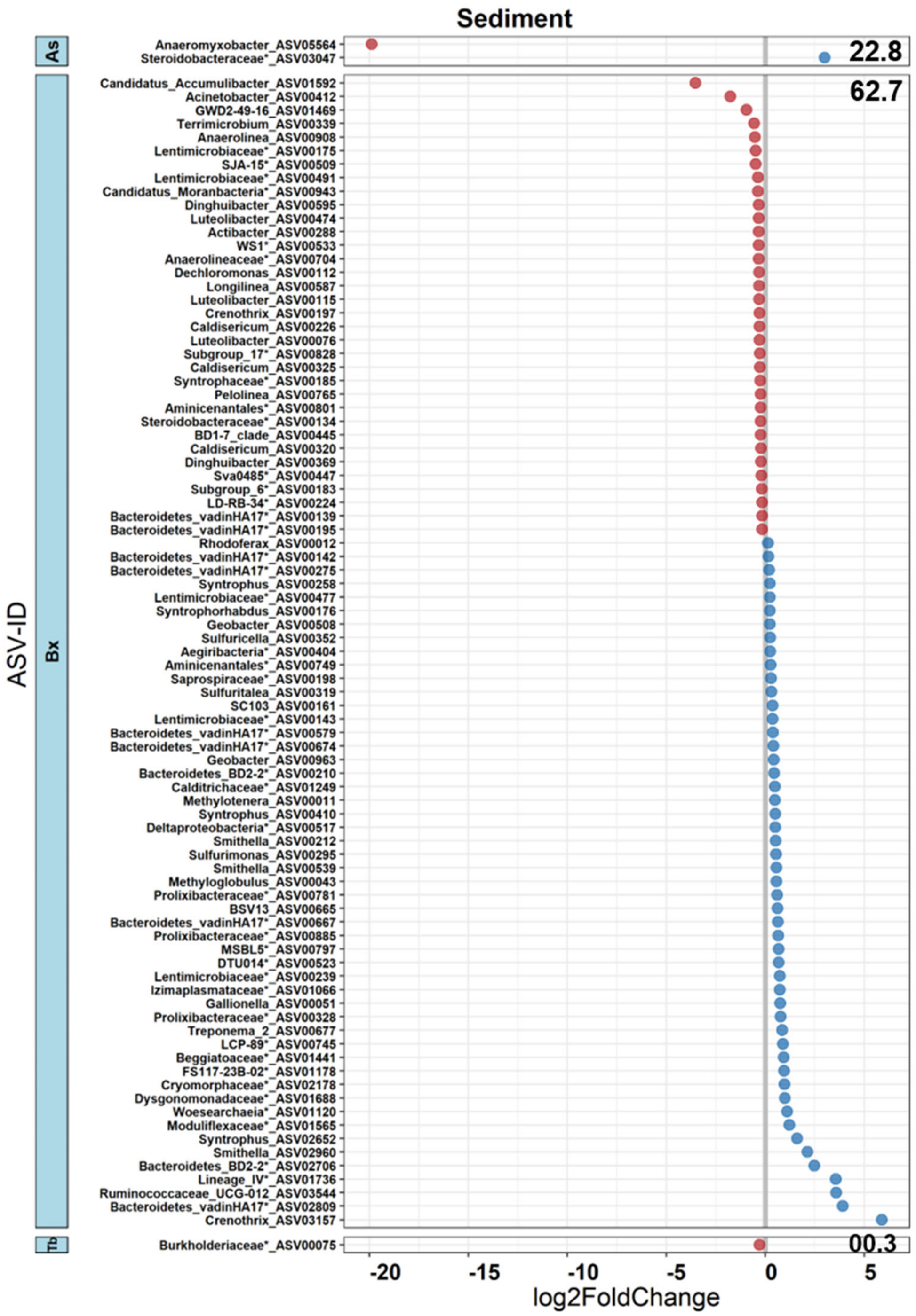
Sediment
As
Anaeromyxobacter_ASV05564
Steroidobacteraceae*_ASV03047
22.8
Bx
Candidatus_Accumulibacter_ASV01592
Acinetobacter_ASV00412
GWD2-49-16_ASV01469
Terrimicrobium_ASV00339
Anaerolinea_ASV00908
Lentimicrobiaceae*_ASV00175
SJA-15*_ASV00509
Lentimicrobiaceae*_ASV00491
Candidatus_Moranbacteria*_ASV00943
Dinghuibacter_ASV00595
Luteolibacter_ASV00474
Actibacter_ASV00288
WS1*_ASV00533
Anaerolineaceae*_ASV00704
Dechloromonas_ASV00112
Longilinea_ASV00587
Luteolibacter_ASV00115
Crenothrix_ASV00197
Caldisericum_ASV00226
Luteolibacter_ASV00076
Subgroup_17*_ASV00828
Caldisericum_ASV00325
Syntrophaceae*_ASV00185
Pelolinea_ASV00765
Aminicenantales*_ASV00801
Steroidobacteraceae*_ASV00134
BD1-7_clade_ASV00445
Caldisericum_ASV00320
Dinghuibacter_ASV00369
Sva0485*_ASV00447
Subgroup_6*_ASV00183
LD-RB-34*_ASV00224
Bacteroidetes_vadinHA17*_ASV00139
Bacteroidetes_vadinHA17*_ASV00195
Rhodoferax_ASV00012
Bacteroidetes_vadinHA17*_ASV00142
Bacteroidetes_vadinHA17*_ASV00275
Syntrophus_ASV00258
Lentimicrobiaceae*_ASV00477
Syntrophorhabdus_ASV00176
Geobacter_ASV00508
Sulfuricella_ASV00352
Aegiribacteria*_ASV00404
Aminicenantales*_ASV00749
Saprospiraceae*_ASV00198
Sulfuritalea_ASV00319
SC103_ASV00161
Lentimicrobiaceae*_ASV00143
Bacteroidetes_vadinHA17*_ASV00579
Bacteroidetes_vadinHA17*_ASV00674
Geobacter_ASV00963
Bacteroidetes_BD2-2*_ASV00210
Calditrichaceae*_ASV01249
Methylotenera_ASV00011
Syntrophus_ASV00410
Deltaproteobacteria*_ASV00517
Smithella_ASV00212
Sulfurimonas_ASV00295
Smithella_ASV00539
Methyloglobulus_ASV00043
Prolixibacteraceae*_ASV00781
BSV13_ASV00665
Bacteroidetes_vadinHA17*_ASV00667
Prolixibacteraceae*_ASV00885
MSBL5*_ASV00797
DTU014*_ASV00523
Lentimicrobiaceae*_ASV00239
Izimaplasmataceae*_ASV01066
Gallionella_ASV00051
Prolixibacteraceae*_ASV00328
Treponema_2_ASV00677
LCP-89*_ASV00745
Beggiatoaceae*_ASV01441
FS117-23B-02*_ASV01178
Cryomorphaceae*_ASV02178
Dysgonomonadaceae*_ASV01688
Woesearchaeia*_ASV01120
Moduliflexaceae*_ASV01565
Syntrophus_ASV02652
Smithella_ASV02960
Bacteroidetes_BD2-2*_ASV02706
Lineage_IV*_ASV01736
Ruminococcaceae_UCG-012_ASV03544
Bacteroidetes_vadinHA17*_ASV02809
Crenothrix_ASV03157
62.7
Th
Burkholderiaceae*_ASV00075
00.3
ASV-ID
-20
-15
-10
-5
0
5
log2FoldChange

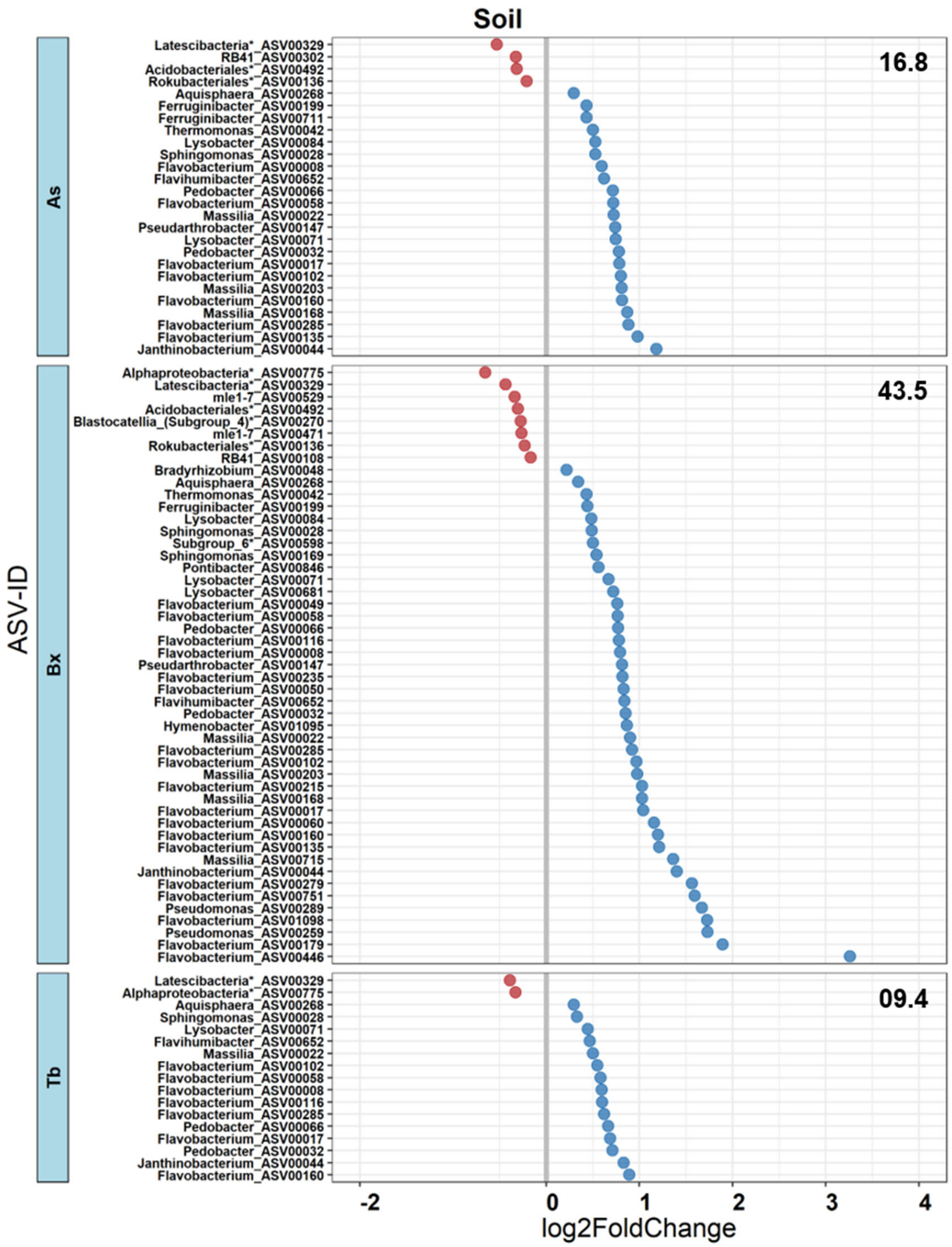

Soil
ASV-ID
As
16.8
Latescibacteria*_ASV00329
RB41_ASV00302
Acidobacteriales*_ASV00492
Rokubacteriales*_ASV00136
Aquisphaera_ASV00268
Ferruginibacter_ASV00199
Ferruginibacter_ASV00711
Thermomonas_ASV00042
Lysobacter_ASV00084
Sphingomonas_ASV00028
Flavobacterium_ASV00008
Flavihumibacter_ASV00652
Pedobacter_ASV00066
Flavobacterium_ASV00058
Massilia_ASV00022
Pseudarthrobacter_ASV00147
Lysobacter_ASV00071
Pedobacter_ASV00032
Flavobacterium_ASV00017
Flavobacterium_ASV00102
Massilia_ASV00203
Flavobacterium_ASV00160
Massilia_ASV00168
Flavobacterium_ASV00285
Flavobacterium_ASV00135
Janthinobacterium_ASV00044
Bx
43.5
Alphaproteobacteria*_ASV00775
Latescibacteria*_ASV00329
mle1-7_ASV00529
Acidobacteriales*_ASV00492
Blastocatellia_(Subgroup_4)*_ASV00270
mle1-7_ASV00471
Rokubacteriales*_ASV00136
RB41_ASV00108
Bradyrhizobium_ASV00048
Aquisphaera_ASV00268
Thermomonas_ASV00042
Ferruginibacter_ASV00199
Lysobacter_ASV00084
Sphingomonas_ASV00028
Subgroup_6*_ASV00598
Sphingomonas_ASV00169
Pontibacter_ASV00846
Lysobacter_ASV00071
Lysobacter_ASV00681
Flavobacterium_ASV00049
Flavobacterium_ASV00058
Pedobacter_ASV00066
Flavobacterium_ASV00116
Flavobacterium_ASV00008
Pseudarthrobacter_ASV00147
Flavobacterium_ASV00235
Flavobacterium_ASV00050
Flavihumibacter_ASV00652
Pedobacter_ASV00032
Hymenobacter_ASV01095
Massilia_ASV00022
Flavobacterium_ASV00285
Flavobacterium_ASV00102
Massilia_ASV00203
Flavobacterium_ASV00215
Massilia_ASV00168
Flavobacterium_ASV00017
Flavobacterium_ASV00060
Flavobacterium_ASV00160
Flavobacterium_ASV00135
Massilia_ASV00715
Janthinobacterium_ASV00044
Flavobacterium_ASV00279
Flavobacterium_ASV00751
Pseudomonas_ASV00289
Flavobacterium_ASV01098
Pseudomonas_ASV00259
Flavobacterium_ASV00179
Flavobacterium_ASV00446
Tb
09.4
Latescibacteria*_ASV00329
Alphaproteobacteria*_ASV00775
Aquisphaera_ASV00268
Sphingomonas_ASV00028
Lysobacter_ASV00071
Flavihumibacter_ASV00652
Massilia_ASV00022
Flavobacterium_ASV00102
Flavobacterium_ASV00058
Flavobacterium_ASV00008
Flavobacterium_ASV00116
Flavobacterium_ASV00285
Pedobacter_ASV00066
Flavobacterium_ASV00017
Pedobacter_ASV00032
Janthinobacterium_ASV00044
Flavobacterium_ASV00160
-2
0
1
2
3
4
log2FoldChange

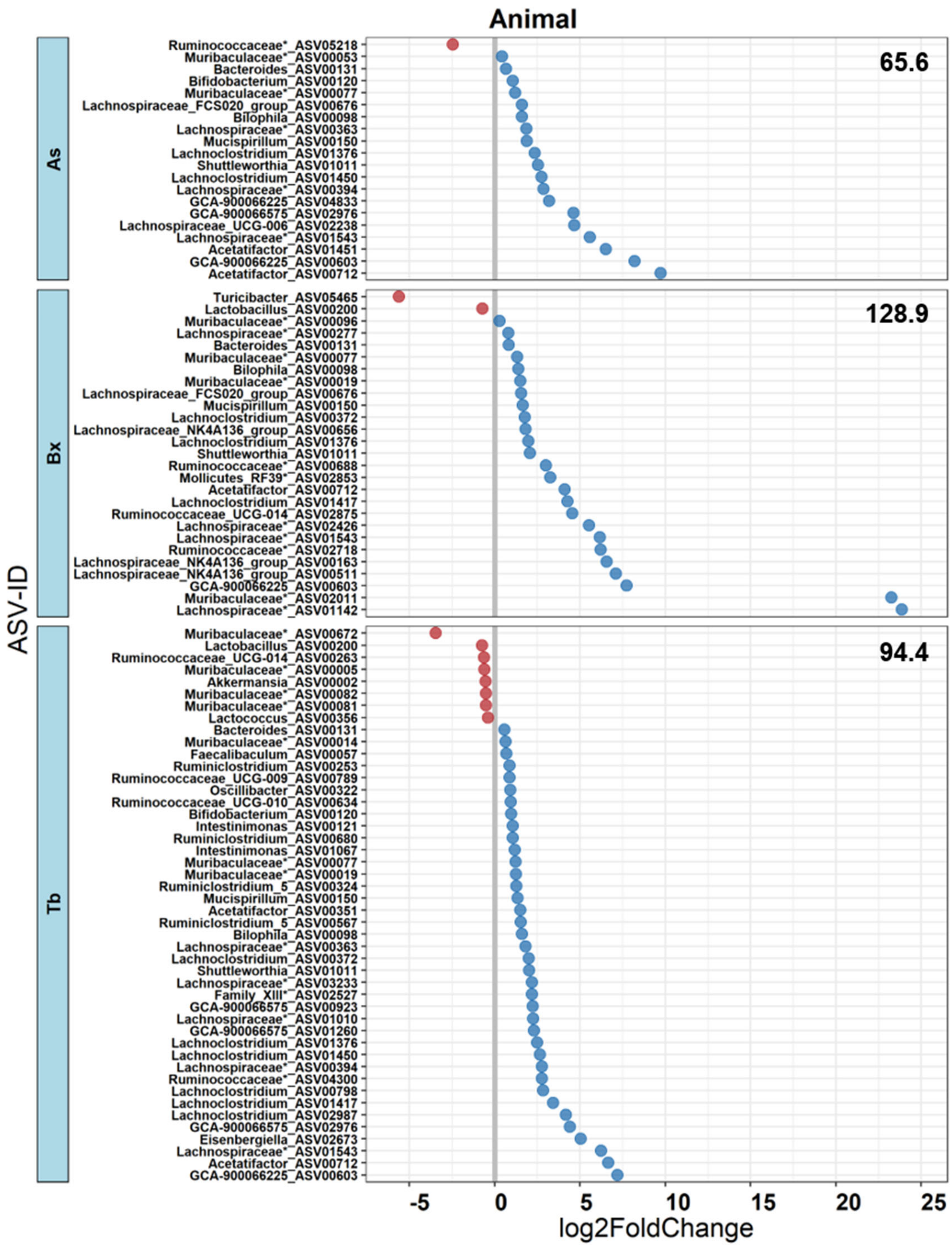


**Figure S7 | Stress-sensitive ASVs and Influence Scores**
The Figure displays for each component the ASVs that differed significantly ($P \leq 0.05$) in mean relative abundance due to the different chemical treatments As, Bx and Tb relative to their controls. Genus taxonomy is provided next to the ASV-ID. The statistics outputs of the negative binomial-based Wald tests are documented in the **Database S1**. Decrease and increase of log2 fold changes are colored in red and blue, respectively. For each microbiome and stress combination, an Influence Score (indicated in bold numbers, top right) was calculated. The Influence Score considers both the number of affected ASVs and their log-fold change (see methods).

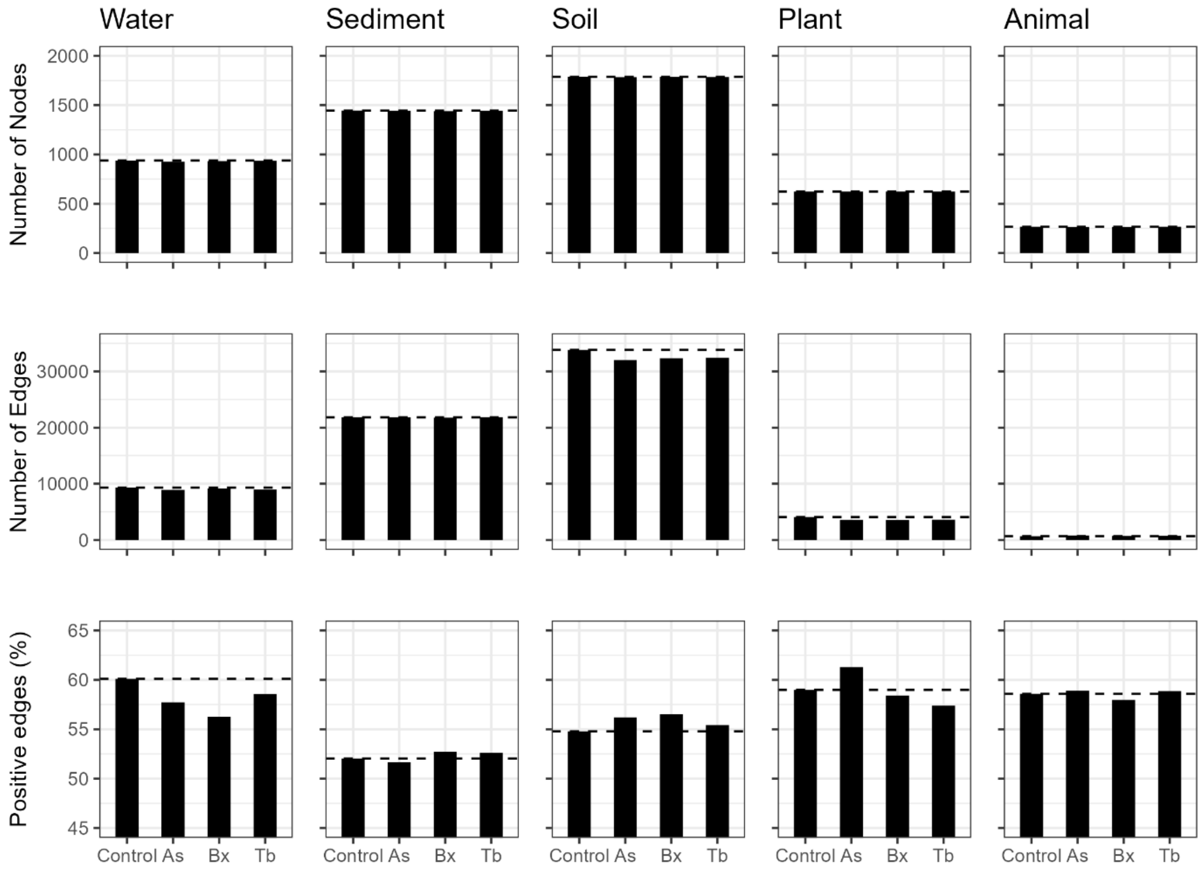


**Figure S8 | Network characteristics**
Co-occurrence networks were generated for each treatment (control, As, Bx and Tb treated) and all microbiomes. Then the canonical parameters extracted from these networks. Nodes represent ASVs and edges connect nodes that correlate (positive or negatively) in their abundances. Number of nodes, numbers of edges and the percentage of positive edges are displayed here while additional network characteristics such as node degree, betweenness centrality, closeness centrality, transitivity are listed in **Table S7**.

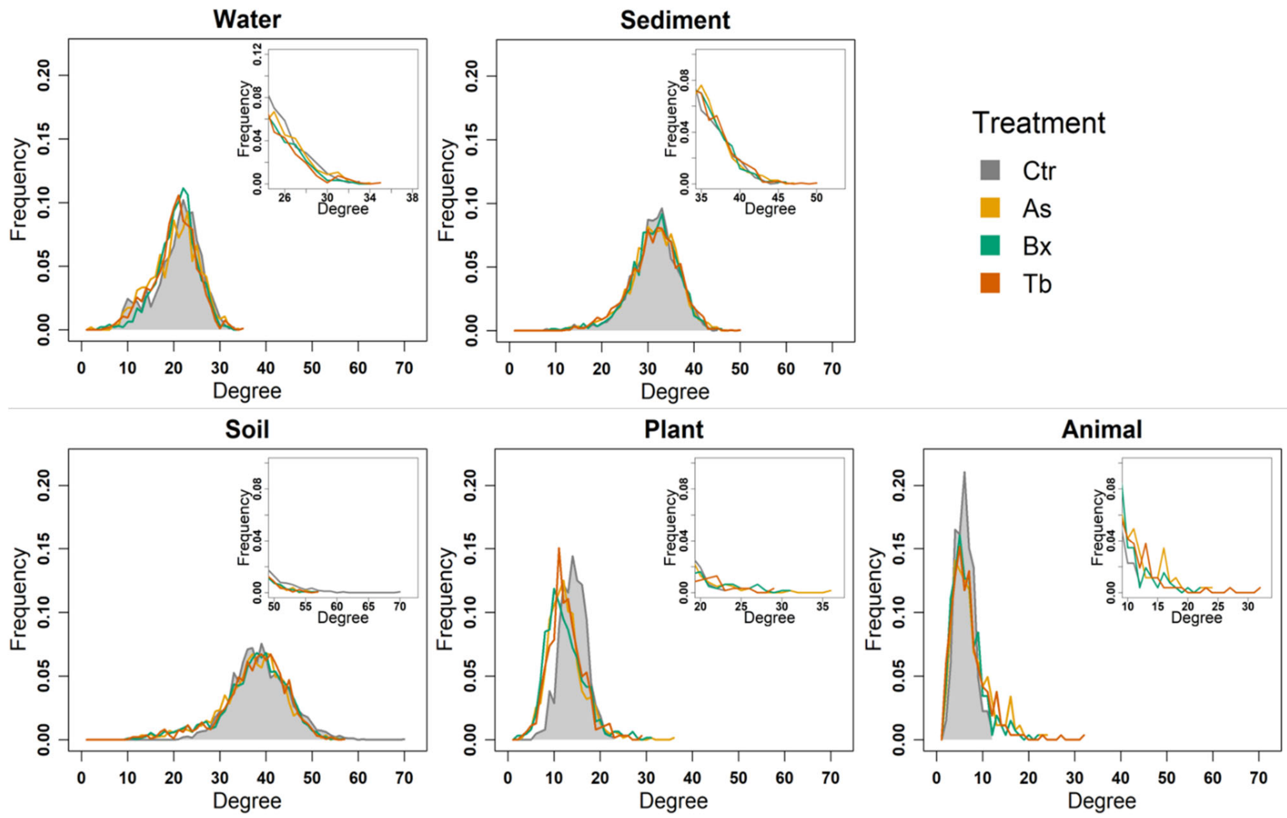


**Figure S9 | Network degree distributions**
For each component, the frequency distribution of the node degrees in each microbiome network (control, As, Bx, and Tb treated) is displayed. The insets highlight the distributions in the regions of higher node degrees to better visualize the differences for the nodes having high degrees and a tail exhibiting significantly higher connections.

#### Supplementary Tables

**Table S1 | Basic parameters of the experimental soil**

Analyses of soil pH, proportions of physical particles, micro-nutrient, macro-nutrient and elements were performed for a basic characterization of the experimental soil (n=4, s.e.m.).

| Measurements | Unit | Mean | ± SEM |
|---|---|---|---|
| pH | - | 6.61 | ± 0.04 |
| Clay | % | 10.2 | ± 0.8 |
| Silt | % | 53 | ± 2 |
| Sand | % | 37 | ± 3 |
| Plant available phosphorous | (mg/kg) | 2.41 | ± 0.02 |
| Total carbon | (g/kg) | 26.49 | ± 0.07 |
| Total organic carbon | (g/kg) | 25.41 | ± 0.09 |
| Nitrogen | (g/kg) | 2.91 | ± 0.01 |
| Sulfur | (g/kg) | 0.35 | ± 0.03 |
| Arsenic | (mg/kg) | 2.9 | ± 0.5 |
| Magnesium | (g/kg) | 4.1 | ± 0.5 |
| Potassium | (g/kg) | 1.5 | ± 0.1 |
| Aluminium | (g/kg) | 10.8 | ± 0.8 |
| Iron | (g/kg) | 17.7 | ± 0.7 |
| Manganese | (g/kg) | 0.74 | ± 0.02 |
| Chromium | (mg/kg) | 23 | ± 2 |
| Copper | (mg/kg) | 20 | ± 3 |
| Zinc | (mg/kg) | 66 | ± 3 |
| Cadmium | (mg/kg) | 0.18 | ± 0.00 |
| Lead | (mg/kg) | 18.5 | ± 0.7 |

**Table S2 | Factors affecting beta diversity**

PERMANOVA based on Bray-Curtis metrics was used to test for significant effects of treatment, timepoint, concentration and sample type and interactions thereof based on 1000 permutations. Significant *P*-values are highlighted in bold.

| **Variables** | **SumsOfSq** | **MeanSq** | **F.Model** | $R^2$ | ***P*** |
|---|---|---|---|---|---|
| Treatment | 3.03 | 1.01 | 15.03 | 0.007 | **0.001** |
| Timepoint | 22.78 | 11.39 | 169.66 | 0.056 | **0.001** |
| Concentration | 9.62 | 3.21 | 47.75 | 0.024 | **0.001** |
| Sampletype | 291.53 | 72.88 | 1085.67 | 0.718 | **0.001** |
| Library | 0.42 | 0.10 | 1.55 | 0.001 | 0.059 |
| Treatment:Timepoint | 0.62 | 0.12 | 1.84 | 0.002 | **0.011** |
| Treatment:Concentration | 0.45 | 0.11 | 1.69 | 0.001 | **0.030** |
| Timepoint:Concentration | 0.19 | 0.10 | 1.45 | 0.000 | 0.133 |
| Treatment:Sampletype | 2.89 | 0.24 | 3.59 | 0.007 | **0.001** |
| Timepoint:Sampletype | 5.01 | 0.84 | 12.44 | 0.012 | **0.001** |
| Concentration:Sampletype | 0.76 | 0.09 | 1.41 | 0.002 | **0.047** |
| Treatment:Timepoint:Concentration | 0.35 | 0.09 | 1.30 | 0.001 | 0.164 |
| Treatment:Timepoint:Sampletype | 1.20 | 0.10 | 1.49 | 0.003 | **0.009** |
| Treatment:Concentration:Sampletype | 1.61 | 0.10 | 1.5 | 0.004 | **0.002** |
| Timepoint:Concentration:Sampletype | 0.72 | 0.09 | 1.33 | 0.002 | 0.063 |
| Treatment:Timepoint:Conc:Sampletype | 1.4 | 0.09 | 1.37 | 0.149 | **0.040** |

**Table S3 | Factors affecting alpha diversity**
Factorial ANOVA was used to assess ASV richness and Shannon diversity for significant effects of treatment, timepoint, concentration and sample type and interactions thereof based on 1000 permutations. Significant *P*-values are highlighted in bold.

| Variables | SumsOfSq | MeanSq | F.Model | *P* |
|---|---|---|---|---|
| **Richness (number of observed ASVs)** | | | | |
| Treatment | 650875 | 216958 | 30.32 | <**2.00E-16** |
| Timepoint | 9906990 | 4953495 | 692.14 | <**2.00E-16** |
| Concentration | 1885926 | 628642 | 87.84 | <**2.00E-16** |
| Sampletype | 72660876 | 18165219 | 2538.19 | <**2.00E-16** |
| Library | 1015927 | 253982 | 35.49 | <**2.00E-16** |
| Treatment:Timepoint | 15902 | 3180 | 0.44 | 0.818 |
| Treatment:Concentration | 128359 | 32090 | 4.48 | **0.001** |
| Timepoint:Concentration | 82984 | 41492 | 5.80 | **0.003** |
| Treatment:Sampletype | 403823 | 33652 | 4.70 | **1.74E-07** |
| Timepoint:Sampletype | 485426 | 80904 | 11.31 | **3.19E-12** |
| Concentration:Sampletype | 45589 | 5699 | 0.80 | 0.606 |
| Treatment:Timepoint:Concentration | 44863 | 11216 | 1.57 | 0.181 |
| Treatment:Timepoint:Sampletype | 271081 | 22590 | 3.16 | **0.000** |
| Treatment:Concentration:Sampletype | 491577 | 30724 | 4.29 | **3.61E-08** |
| Timepoint:Concentration:Sampletype | 196440 | 24555 | 3.43 | **0.001** |
| Treatment:Timepoint:Conc:Sampletype | 399824 | 24989 | 3.49 | **4.02E-06** |
| **Shannon diversity** | | | | |
| Treatment | 7 | 2.35 | 37.24 | <**2.00E-16** |
| Timepoint | 156.6 | 78.31 | 1242.07 | <**2.00E-16** |
| Concentration | 37.9 | 12.64 | 200.48 | <**2.00E-16** |
| Sampletype | 939.9 | 234.99 | 3726.93 | <**2.00E-16** |
| Library | 0.8 | 0.2 | 3.16 | **0.014** |
| Treatment:Timepoint | 0.4 | 0.09 | 1.39 | 0.225 |
| Treatment:Concentration | 0.9 | 0.22 | 3.55 | **0.007** |
| Timepoint:Concentration | 0.8 | 0.41 | 6.43 | **0.002** |
| Treatment:Sampletype | 8 | 0.67 | 10.58 | <**2.00E-16** |
| Timepoint:Sampletype | 6.2 | 1.04 | 16.52 | <**2.00E-16** |
| Concentration:Sampletype | 1 | 0.13 | 2.03 | **0.040** |
| Treatment:Timepoint:Concentration | 0.4 | 0.09 | 1.45 | 0.215 |
| Treatment:Timepoint:Sampletype | 1.2 | 0.1 | 1.60 | 0.087 |
| Treatment:Concentration:Sampletype | 2.5 | 0.16 | 2.47 | **0.001** |
| Timepoint:Concentration:Sampletype | 0.6 | 0.07 | 1.10 | 0.361 |
| Treatment:Timepoint:Conc:Sampletype | 1.4 | 0.09 | 1.37 | 0.149 |

**Table S4 | Pairwise factors affecting beta diversity**
Pairwise PERMANOVAs tests were performed on the Bray-Curtis metrics to test for significant differences between treatments and their respective controls. Significant *P*-values are highlighted in bold.

| Pairs | SumsOfSq | F.Model | $R^2$ | *P*.value | *P*.adjusted |
|---|---|---|---|---|---|
| Ctr.water vs As.water | 0.101 | 1.577 | 0.024 | 0.085 | 0.085 |
| Ctr.water vs Bx.water | 0.138 | 2.220 | 0.034 | 0.032 | **0.033** |
| Ctr.water vs Tb.water | 0.578 | 9.030 | 0.120 | 0.001 | **0.001** |
| Ctr.sediment vs As.sediment | 0.072 | 1.265 | 0.018 | 0.045 | **0.046** |
| Ctr.sediment vs Bx.sediment | 0.217 | 3.868 | 0.052 | 0.001 | **0.001** |
| Ctr.sediment vs Tb.sediment | 0.092 | 1.541 | 0.022 | 0.005 | **0.005** |
| Ctr.soil vs As.soil | 0.349 | 5.017 | 0.043 | 0.001 | **0.001** |
| Ctr.soil vs Bx.soil | 0.515 | 7.388 | 0.062 | 0.001 | **0.001** |
| Ctr.soil vs Tb.soil | 0.271 | 4.168 | 0.037 | 0.001 | **0.001** |
| Ctr.plant vs As.plant | 0.232 | 2.255 | 0.021 | 0.010 | **0.010** |
| Ctr.plant vs Bx.plant | 0.237 | 2.263 | 0.021 | 0.016 | **0.016** |
| Ctr.plant vs Tb.plant | 0.238 | 2.378 | 0.023 | 0.009 | **0.009** |
| Ctr.animal vs As.animal | 0.286 | 3.418 | 0.032 | 0.004 | **0.004** |
| Ctr.animal vs Bx.animal | 0.220 | 2.834 | 0.028 | 0.004 | **0.004** |
| Ctr.animal vs Tb.animal | 0.347 | 5.200 | 0.047 | 0.001 | **0.001** |

**Table S5 | Constrained ordination for treatment effect in each component**
We performed constrained ordination of Bray-Curtis distances using R function *capscale* based on 999 permutations and assess the significance of constraints using ANOVA like permutation test. Significant *P*-values are highlighted in bold.

| Variable | Df | SumsOfSq | F.Model | *P* |
|---|---|---|---|---|
| Water | 3 | 1.254 | 6.995 | **0.001** |
| Sediment | 3 | 0.451 | 2.582 | **0.001** |
| Soil | 3 | 0.689 | 3.441 | **0.001** |
| Plant | 3 | 0.563 | 1.838 | **0.002** |
| Animal | 3 | 0.812 | 2.716 | **0.001** |

**Table S6 | Testing homogeneity of group dispersion**
We tested homogeneity of group dispersions of Bray-Curtis distances using PERMDISP tests of the R function *betadisper* based on 999 permutations. Significant *P*-values are highlighted in bold.

| Variable | Df | SumsOfSq | MeanSq | F.Model | *P* |
|---|---|---|---|---|---|
| Water-Treatment | 3 | 0.024 | 0.008 | 1.247 | 0.283 |
| Sediment-Treatment | 3 | 0.003 | 0.001 | 1.684 | 0.158 |
| Soil-Treatment | 3 | 0.015 | 0.005 | 1.837 | 0.137 |
| Plant-Treatment | 3 | 0.013 | 0.004 | 2.511 | **0.048** |
| Animal-Treatment | 3 | 0.107 | 0.036 | 6.491 | **0.001** |

**Table S7 | Network properties**

Co-occurrence networks were generated for all microbiomes and each treatment (control, As, Bx and Tb). Canonical network parameters were extracted from these networks. Hub nodes (*) were selected based on hub centrality score value ≥0.7. Significant differences (*P*≤0.001) based on 10,000 bootstrap replicates are marked in bold.

| Source | Network Properties | Control | As | Bx | Tb |
|---|---|---|---|---|---|
| **Water** | Number of nodes | 939 | 927 | 932 | 936 |
| | Edges-pos | 5582 | 5135 | 5158 | 5249 |
| | Edges-neg | 3706 | 3762 | 4008 | 3714 |
| | Edges per node | 8.9 | 8.34 | 8.62 | 8.74 |
| | Mean Degree | 19.818 | **19.142** | **19.711** | **19.179** |
| | Mean Betweenness | 18393.7x10$^{-7}$ | **18728.3x10$^{-7}$** | **17974.1x10$^{-7}$** | **18417.6x10$^{-7}$** |
| | Mean Closeness | 3698104.0x10$^{-7}$ | **3664606.0x10$^{-7}$** | **3760383.0x10$^{-7}$** | **3687542.0x10$^{-7}$** |
| | Mean Transitivity | 709553.0x10$^{-7}$ | **719553.6x10$^{-7}$** | **660084.1x10$^{-7}$** | **647006.4x10$^{-7}$** |
| | Hub Nodes* | 112 | 162 | 81 | 45 |
| | Best Fit Dist | Undecided | Undecided | Undecided | Undecided |
| **Sediment** | Number of nodes | 1445 | 1444 | 1439 | 1444 |
| | Edges-pos | 11366 | 11281 | 11486 | 11502 |
| | Edges-neg | 10479 | 10559 | 10295 | 10356 |
| | Edges per node | 9.23 | 8.63 | 8.35 | 8.52 |
| | Mean Degree | 30.216 | **30.289** | **30.342** | **30.379** |
| | Mean Betweenness | 10518.5x10$^{-7}$ | **10569.6x10$^{-7}$** | **10625.1x10$^{-7}$** | **10637.8x10$^{-7}$** |
| | Mean Closeness | 3969524.0x10$^{-7}$ | **3970372.0x10$^{-7}$** | **3970332.0x10$^{-7}$** | **3968447.0x10$^{-7}$** |
| | Mean Transitivity | 507293.0x10$^{-7}$ | **531003.9x10$^{-7}$** | **530470.9x10$^{-7}$** | **542015.7x10$^{-7}$** |
| | Hub Nodes* | 621 | 506 | 423 | 323 |
| | Best Fit Dist | Undecided | Undecided | Undecided | Undecided |
| **Soil** | Number of nodes | 1787 | 1784 | 1786 | 1785 |
| | Edges-pos | 18534 | 17993 | 18272 | 17967 |
| | Edges-neg | 15296 | 14027 | 14051 | 14453 |
| | Edges per node | 18.93 | 17.94 | 18.09 | 18.16 |
| | Mean Degree | 37.850 | **35.926** | **36.123** | **36.283** |
| | Mean Betweenness | 8200.4x10$^{-7}$ | **8281.4x10$^{-7}$** | **8202.6x10$^{-7}$** | **8201.2x10$^{-7}$** |
| | Mean Closeness | 4060475.0x10$^{-7}$ | **4049828.0x10$^{-7}$** | **4058185.0x10$^{-7}$** | **4064149.0x10$^{-7}$** |
| | Mean Transitivity | 669839.9x10$^{-7}$ | **437038.3x10$^{-7}$** | **431221.5x10$^{-7}$** | **428233.6x10$^{-7}$** |
| | Hub Nodes* | 78 | 490 | 464 | 716 |
| | Best Fit Dist | Power Law | Undecided | Power Law | Undecided |
| **Plant** | Number of nodes | 624 | 624 | 624 | 624 |
| | Edges-pos | 2412 | 2182 | 2065 | 2068 |
| | Edges-neg | 1677 | 1378 | 1470 | 1535 |
| | Edges per node | 6.55 | 5.7 | 5.66 | 5.77 |
| | Mean Degree | 13.149 | **11.371** | **11.321** | **11.502** |
| | Mean Betweenness | 29529.8x10$^{-7}$ | **31395.3x10$^{-7}$** | **31630.2x10$^{-7}$** | **30876.7x10$^{-7}$** |
| | Mean Closeness | 3547761.0x10$^{-7}$ | **3386832.0x10$^{-7}$** | **3386474.0x10$^{-7}$** | **3418976.0x10$^{-7}$** |
| | Mean Transitivity | 680524.6x10$^{-7}$ | **568188.9x10$^{-7}$** | **503979.0x10$^{-7}$** | **558959.2x10$^{-7}$** |
| | Hub Nodes* | 74 | 2 | 11 | 8 |
| | Best Fit Dist | Undecided | Undecided | Log Normal | Undecided |
| **Animal** | Number of nodes | 266 | 264 | 264 | 266 |
| | Edges-pos | 396 | 486 | 422 | 455 |
| | Edges-neg | 280 | 339 | 306 | 318 |
| | Edges per node | 2.54 | 2.79 | 2.75 | 2.9 |
| | Mean Degree | 5.089 | **6.263** | **5.584** | **5.810** |
| | Mean Betweenness | 103175.2x10$^{-7}$ | **090283.2x10$^{-7}$** | **099789.8x10$^{-7}$** | **092116.9x10$^{-7}$** |
| | Mean Closeness | 2699416.0x10$^{-7}$ | **3028462.0x10$^{-7}$** | **2825292.0x10$^{-7}$** | **2948858.0x10$^{-7}$** |
| | Mean Transitivity | 686935.8x10$^{-7}$ | **838205.8x10$^{-7}$** | **941475.4x10$^{-7}$** | **694528.4x10$^{-7}$** |
| | Hub Nodes* | 14 | 3 | 3 | 4 |
| | Best Fit Dist | Log Normal | Log Normal | Log Normal | Log Normal |

### Supplementary Data

#### Database S1

For each component, we identified the ASVs that differed significantly in mean relative abundance due to the chemical stressor treatments, by performing differential abundance analysis with DESeq2, i.e. between Ctr-As, Ctr-Bx, Ctr-Tb using negative binomial-based Wald tests ($P \leq 0.05$). The individual ASVs can be retrieved in the Database S1, which consists of a R object that can be easily imported and further used for downstream analyses as follows.

````
```{R}
library(DESeq2)
DatabaseS1 <- readRDS("DatabaseS1.RDS")
names(DatabaseS1)
```
````

```
 [1] "CMRE5_High_mouse_CtrAs1_res"    "CMRE5_High_mouse_CtrBx1_res"
 [3] "CMRE5_High_mouse_CtrTb2_res"    "CMRE5_High_Root_CtrAs2_res"
 [5] "CMRE5_High_Root_CtrBx2_res"     "CMRE5_High_Root_CtrTb2_res"
 [7] "CMRE5_High_Sediment_CtrAs2_res" "CMRE5_High_Sediment_CtrBx2_res"
 [9] "CMRE5_High_Sediment_CtrTb2_res" "CMRE5_High_Soil_CtrAs2_res"
[11] "CMRE5_High_Soil_CtrBx2_res"     "CMRE5_High_Soil_CtrTb2_res"
[13] "CMRE5_High_Water_CtrAs2_res"    "CMRE5_High_Water_CtrBx2_res"
[15] "CMRE5_High_Water_CtrTb2_res"
```

````
```{R}
DatabaseS1[1]
```
````

```
$CMRE5_High_mouse_CtrAs1_res
log2 fold change (MLE): Treatment As vs Ctr
Wald test p-value: Treatment As vs Ctr
DataFrame with 437 rows and 6 columns
           baseMean log2FoldChange     lfcSE       stat      pvalue        padj
          <numeric>      <numeric> <numeric>  <numeric>   <numeric>   <numeric>
ASV_00603  39.97372        8.21135  0.497260   16.51320 2.94806e-61 1.28830e-58
ASV_00712  26.07167        9.71390  1.489438    6.52185 6.94439e-11 1.51735e-08
ASV_00077 290.12560        1.19942  0.234586    5.11293 3.17207e-07 4.62065e-05
ASV_02976   3.31713        4.61929  1.050607    4.39678 1.09867e-05 1.20030e-03
ASV_01011  23.56870        2.53288  0.597790    4.23708 2.26450e-05 1.97917e-03
...             ...            ...       ...        ...         ...         ...
ASV_01672 10.373972      0.0132655  0.600272  0.0220991    0.982369    0.991123
ASV_03503  2.126923     -0.0195988  1.262301 -0.0155262    0.987612    0.991123
ASV_05638  0.583347     -0.0280265  2.189043 -0.0128031    0.989785    0.991123
ASV_03947  1.389272     -0.0378086  2.802188 -0.0134925    0.989235    0.991123
ASV_04018  1.271429     -0.0191505  1.721193 -0.0111263    0.991123    0.991123
```
